# Hidden genetic variation in plasticity provides the potential for rapid adaptation to novel environments

**DOI:** 10.1101/2020.10.26.356451

**Authors:** Greg M. Walter, James Clark, Delia Terranova, Salvatore Cozzolino, Antonia Cristaudo, Simon J. Hiscock, Jon Bridle

## Abstract

Rapid environmental change is forcing populations into novel environments where plasticity will no longer maintain fitness. When populations are exposed to novel environments, evolutionary theory predicts that genetic variation in fitness will increase and should be associated with genetic differences in plasticity. If true, then genetic variation in plasticity can increase adaptive potential in novel environments, and population persistence via rapid adaptation is more likely. To test whether genetic variation in fitness increases in novel environments and is associated with plasticity, we transplanted 8,149 clones of 314 genotypes of a Sicilian daisy (*Senecio chrysanthemifolius*) within and outside its native range, and quantified genetic variation in fitness, and plasticity in leaf traits and gene expression. Although mean fitness declined by 87% in the novel environment, genetic variance in fitness increased threefold and was correlated with plasticity in leaf traits. High fitness genotypes showed greater plasticity in gene expression, but lower plasticity in most leaf traits. Interestingly, genotypes with greater fitness in the novel environment had the lowest fitness at the native site. These results suggest that standing genetic variation in plasticity could help populations to persist and adapt to novel environments, despite remaining hidden in native environments.

## INTRODUCTION

Understanding how populations and ecological communities will respond to rapid environmental change remains a fundamental challenge (Bridle and Hoffmann 2022; Parmesan 2006; Shaw and Etterson 2012). Populations respond to new environments either by genotypes adjusting their phenotypes to match changing conditions (adaptive plasticity) (Charmantier et al. 2008; Via et al. 1995), or by increases in the frequency of beneficial alleles that increase fitness and promote adaptation (termed ‘evolutionary rescue’) (Bell and Gonzalez 2009; Gomulkiewicz and Holt 1995). However, if evolutionary rescue relies on alleles that increase fitness via beneficial plastic responses, then genetic variation in plasticity will be critical for persistence in novel environments (Chevin and Hoffmann 2017; Chevin and Lande 2011; Kelly 2019; Lande 2009).

Given that plasticity can only evolve to match the environmental variation previously encountered within the native range of a species, existing plasticity should only maintain fitness across a species’ current or historical range (Chevin et al. 2013; Ghalambor et al. 2007). Such adaptive plasticity is expected to reduce genetic variation in fitness because plasticity common to all genotypes can successfully maintain fitness in familiar environments (Bradshaw 1991). However, as adaptive plasticity becomes less effective in more novel environments and absolute fitness of the population declines, genetic variation in fitness is expected to increase because genotypes will vary more in their (previously untested) sensitivity to the new conditions (Ashander et al. 2016; Chevin and Lande 2011; Chevin et al. 2010; Hermisson and Wagner 2004; Lande 2009). Such an exposure of genetic variation in novel environments increases the adaptive potential of the population that could allow evolutionary rescue (Agashe et al. 2011; Lande 2009; Nussey et al. 2005), and would suggest that estimates of genetic variation in fitness within a species’ current geographical range will underestimate the adaptive potential of the population when exposed to novel environments.

Populations that experience large reductions in mean absolute fitness 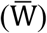 in novel environments will face extinction if they cannot increase 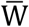 sufficiently quickly to prevent population declines (Hendry et al. 2018; Lande and Shannon 1996; Lynch and Lande 1993). The potential to increase mean fitness in subsequent generations is determined by the ratio of additive genetic variance in fitness to mean fitness 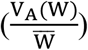 (Bonnet et al. 2019; Fisher 1930; Kulbaba et al. 2019), which determines the potential for evolutionary rescue (Shaw 2019; Sheth et al. 2018). Increased genetic variance in a novel environment will therefore improve the adaptive potential of the population because alleles that can rapidly increase mean fitness are already present.

Although recent studies have found changes in additive genetic variation in fitness within a species range (Kulbaba et al. 2019; Peschel et al. 2020; Sheth et al. 2018), to our knowledge, increased genetic variance in fitness 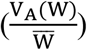 in novel environments has not been identified because studies typically quantify genetic variance in fitness in familiar environments (Hendry et al. 2018) or focus on heritability (Charmantier and Garant 2005; Hoffmann and Merilä 1999), which does not directly quantify adaptive potential (Hansen et al. 2011). Furthermore, studies that link fitness to plasticity in novel environments are even rarer (Steinger et al. 2003; Wang and Althoff 2019). We therefore have a remarkably limited understanding of whether genetic variation in fitness increases in novel environments, or the role of plasticity in explaining any changes in genetic variation in fitness. Without such information, the adaptive potential of populations as they become exposed to novel environments remains largely unknown (Shaw 2019).

We focus on a Sicilian daisy, *Senecio chrysanthemifolius* (Asteraceae), that grows in disturbed habitats at low elevation (c.400-1,000 m.a.s.l [metres above sea level]) on Mount Etna and throughout lowland Sicily (**Fig. 1A**). This species occurs in small patches typically containing fewer than 100 individuals, with patches often separated by 1-2 km. Individuals are occasionally observed at elevations between 1,000 m and 1,500 m (but never above 1,500 m), indicating that these higher elevations represent the edge of the range for *S. chrysanthemifolius*. In natural populations, individuals of *S. chrysanthemifolius* typically live for less than two years and are obligate outcrossers that are pollinated by generalist insects (e.g., hoverflies), with wind-dispersed seeds that can often move hundreds of metres from their parent. Also found on Mt. Etna is a high-elevation sister species, *S. aethnensis*, that is endemic to lava flows above 2,000 m, and is rarely found below 1,500 m. Adaptive divergence between these two *Senecio* species is associated with contrasting leaf morphology and physiology, as well as differences in plasticity (Walter et al. 2022). When reciprocally transplanted in their native habitats, plasticity moved the phenotype of *S. chrysanthemifolius* closer to the native phenotype of *S. aethnensis* at 2,000 m, which was associated with lower mortality than experienced by *S. aethnensis* at low elevations. These data suggest that plasticity observed in *S. chrysanthemifolius* is, to some extent, adaptive at the novel 2,000 m elevation (Walter et al. 2022).

**Fig. 1.**
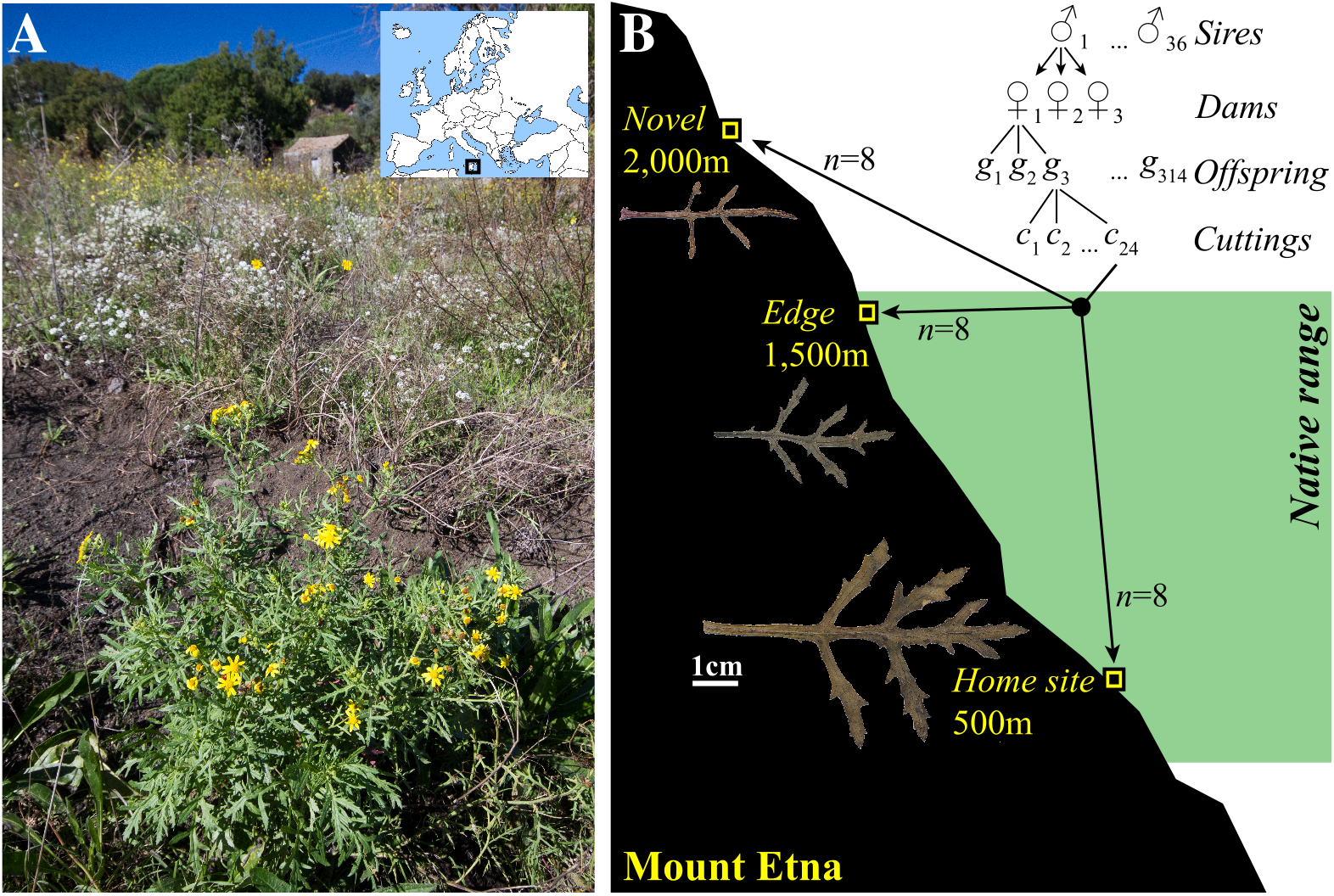
The study system and experimental design. (**A**) *Senecio chrysanthemifolius* in its natural habitat. Inset map presents the location of the study system in relation to Europe. **(B)** Schematic of Mount Etna showing the experimental design. We sampled individuals from five sites in the foothills of Mount Etna, which we crossed in the glasshouse and transplanted cuttings of their offspring at three elevations. We mated 36 sires to 36 dams in 12 blocks of 3×3 (presented in the figure as a nested design for simplicity). Each sire was therefore mated to three dams; we grew three offspring per cross in the glasshouse from which we sampled multiple cuttings that were transplanted at the three elevations. Green shading denotes the native range of *S. chrysanthemifolius* on Mt. Etna and inset leaves show the average change in leaf morphology with elevation observed for a representative genotype.

We collected cuttings from 72 naturally occurring *S. chrysanthemifolius* genotypes on the south-east slopes of Mt. Etna and propagated them in the glasshouse (**Table S1**, **Fig. S1**). We then conducted crosses among them in a paternal half-sibling breeding design (Lynch and Walsh 1998) to produce 314 offspring genotypes, simulating matings that could easily be generated in the natural population (**Fig. 1B**). We then transplanted multiple cuttings (clones) of each offspring genotype at three elevations and quantified fitness, morphology, physiology and gene expression of each genotype in their native environment (500 m), the edge of their range (1,500 m) and a novel environment (2,000 m; **Fig. 1B**).

Using these data, we tested whether adaptive potential increases in novel environments with three predictions: (1) In more familiar environments (500 m and 1,500 m), we would observe high absolute mean fitness 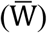 but low additive genetic variance in fitness 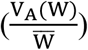 because all genotypes successfully produce appropriate phenotypes within their native range and maintain high fitness. By contrast, at the novel elevation, absolute mean fitness should be reduced and associated with increased genetic variance in fitness that could allow evolutionary rescue to the novel environment (**Fig. 2A**). (2) Significant genetic correlations between trait plasticity and fitness in the novel environment would provide evidence that genotypes with plasticity of a particular magnitude or direction is associated with greater fitness in the novel environment (**Fig. 2B**). (3) Given that changes in gene expression mediate plasticity, genotypes with greater fitness in a novel environment would show differences in their levels of gene expression compared to genotypes with lower fitness, with such changes occurring in genes important for responding to the novel elevation (**Fig. 2C**).

**Fig. 2.**
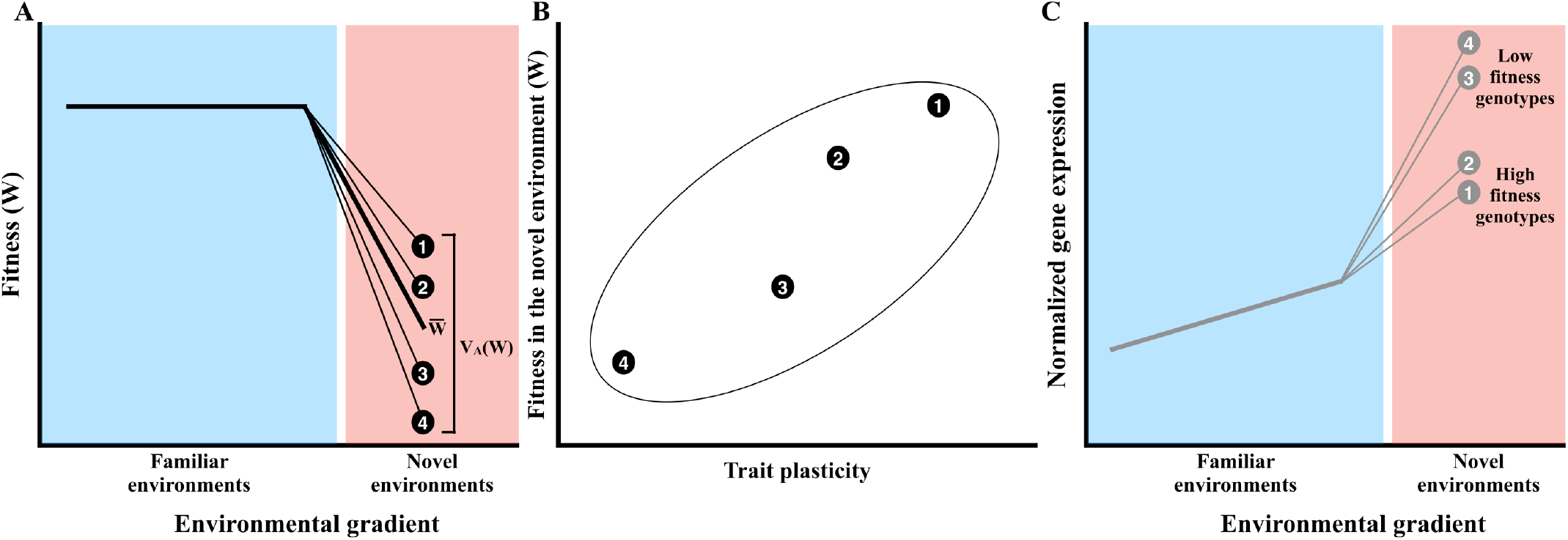
**(A)** Conceptual diagram depicting the predicted change for fitness (W) in response to familiar (blue) and novel (red) environments. The thick line represents the change in mean fitness across environments, and the thin lines with circles represent changes in four different genotypes (represented by different letters) within the population. If all genotypes can effectively buffer familiar environmental variation by generating appropriate phenotypes, we would expect low genetic variance for fitness within the native range to be associated with high mean fitness. This is shown as a high mean fitness in familiar environments, associated with no differences among genotypes. However, in novel environments, we expect a reduction in mean fitness associated with an increase in genetic variance, where differences among genotypes emerge and provide the potential for evolutionary rescue. If genetic differences in plasticity underlie the increase in genetic variation in fitness in the novel environment, we can make two predictions: **(B)** We would expect plasticity to be genetically correlated with genetic variation in fitness. Compared to the genotypes with lower fitness (3-4), the genotypes with greater fitness (1-2) at the novel environment are also associated with different levels of plasticity in a given trait. This creates a moderately strong genetic correlation between trait plasticity and fitness. **(C)** We would also expect that high (1-2) and low (3-4) fitness genotypes to show differences in gene expression. Gray lines represent the gene expression profiles for a single gene (as an example) where gene products are similar for high and low fitness genotypes within the native range, but produce different levels of the gene product in the novel environment. We use gene underexpression as an example, but either under or overexpression could be beneficial in a novel environment. Empirical support for these predictions would suggest that genotypes with particular plastic responses increase fitness in novel environments and provide the potential for evolutionary rescue.

## METHODS

### Genotype sampling and crossing design

To establish the parental generation, in June 2017 we collected cuttings from 72 individuals from five sites <5 km apart on the foothills of Mt. Etna (**Table S1** and **Fig. S1**). The proximity of these sites and the fact that *S. chrysanthemifolius* is insect pollinated and its seeds wind-dispersed mean that gene flow is likely to routinely occur between them. Where possible, we sampled individuals that were at least ten metres apart to minimise chances of sampling close relatives. We removed all branches from mature plants that possessed vegetative material, which we cut into 4-5 cm segments at the glasshouse (Giarre, Italy), dipped them in a rooting plant growth regulator for softwood cuttings (Germon^®^ Bew., Der. NAA 0.5%, L. Gobbi, Italy) and placed each cutting in one cell of an 84-cell tray containing a compressed mix of 1:1 perlite and coconut coir. For three weeks, we kept cuttings in plastic tunnels to maintain humidity and encourage root growth. We then placed one randomly selected cutting per individual in a 30 cm diameter pot with standard potting mix, which we watered regularly. To encourage growth, we suspended 25W LED tubes (TSA Technology, Italy) 1 m above the bench. Once plants produced buds, we covered flowering branches with perforated bread bags to prevent pollinators from entering while allowing airflow. We randomly designated each individual as a dam or sire and grouped them into 12 blocks, each containing three sires (*n*=36) and three dams (*n*=36) (**Table S1**). Because this species is self-incompatible we could mate individuals by removing the flowers from sires and rubbing them on flowers of the dams. Within each block we mated all sires to all dams to produce nine full-sibling families per block (*n*=104 total full-sibling families, with four crosses failing to produce seeds).

Six seeds from each family were germinated by cutting the top off each seed (<1mm) and placing them on moistened filter paper in a plastic Petri dish. Petri dishes were kept in the dark for two days, and then transferred to growth cabinets maintained at 22°C with a 12h: 12h light:dark photoperiod. After one-week we transferred seedlings to the glasshouse where three individuals from each family were grown in 14 cm diameter pots containing standard potting mix (*n=*312 individuals). When the main stem of each seedling reached c.12 cm of growth, we cut the main stem 4 cm above ground level to promote lateral branching and generate enough cuttings for the field transplant.

### Field transplant of the cultivated offspring as cuttings

When branches started producing buds, we took cuttings from all 312 individuals (hereafter, genotypes). Branches were cut into smaller segments each 4-5 cm long with 2-3 leaf nodes. For almost all genotypes we were able to take 21 cuttings (transplant 1). To produce a second round of cuttings from the same genotypes, they were left to regrow for three weeks, at which point we repeated the process to generate another 14 cuttings from each genotype (transplant 2). Due to the deaths of two genotypes in the glasshouse after taking the initial cuttings, we replaced these genotypes with siblings for the second round of cuttings, which led to two extra genotypes (*n*=314 rather than 312). Cuttings were stored in high-humidity tunnels for two weeks until they produced roots and were ready to transplant.

We transplanted 7-10 cuttings from each of the 314 genotypes at each of three transplant sites along an elevational gradient (*n*=c.2,700 cuttings/site; total *N*=8,149 cuttings) that represented typical conditions within the species’ native range (500 m.a.s.l), the edge of its range (1,500 m.a.s.l), and a novel elevation (2,000 m.a.s.l). The 500 m site was located on abandoned land near a local population, the 1,500 m site in an apple and pear orchard where vagrant *S. chrysanthemifolius* individuals were found, and the novel 2,000 m site on a lava flow from c.1980 (**Fig. S1**). Elevations under 1,000 m experience high temperatures that regularly exceed 40°C during summer, and temperatures above 1,000 m frequently drop below 0°C during winter. Soil is characterised as a silty sand at 500 and 1,500 m, but changes to volcanic sand at 2,000 m. Soil chemistry changes gradually across elevation, with lower nutrient content present at higher elevations (Walter et al. 2022).

To prepare experimental plots, the soil surface was cleared of plants and debris, and the soil turned (30 cm deep) immediately prior to transplanting. We transplanted cuttings into large grids of different sizes, spacing each plant 30 cm apart. Transplanted cuttings were maintained moist until they established, after which we reduced watering to minimal amounts that were sufficient to maintain survival for all genotypes while having a minimal effect on growth or flower production.

We transplanted the cuttings in two temporal blocks:

*<u>Transplant 1, 23-25 May 2018:</u>* At each transplant elevation, we randomised six cuttings of each of the 312 genotypes into two replicate environmental blocks (*n*=936 cuttings/block; *n*=1,872 cuttings/transplant site; total N=5,634 cuttings).
*<u>Transplant 2, 10-11 July 2018:</u>* We used all remaining cuttings to transplant two additional cuttings/genotype at each elevation into two additional blocks, as well as replace cuttings lost due to c.10% mortality at 500 m and 2,000 m due to a short spell of intense heat in late June (*n*=321-525 cuttings/block; *n*=718-927 cuttings/transplant site; total N=2,515 cuttings).

### Data collection

Mortality of established cuttings was low, even at the novel elevation (12% across all elevations), allowing us to assay fitness for 6-8 cuttings per genotype at each transplant site. After 5 months of growth (September-October), we counted all flower heads produced by each plant, which we used as an estimate of fitness for each clone. This trait is used routinely to assay fitness in short-lived perennials as total reproductive output for each plant in their first season (e.g. Pujol et al. 2014). Given that plants of *S. chrysanthemifolius* rarely flower more than once, and that the number of flowers was closely associated with the total seeds produced per plant in a previous experiment (Methods S1; Walter et al. 2022), the total number of flowers represents a good proxy of fitness for *S. chrysanthemifolius*.

We measured leaf morphology and pigment content by sampling 3-4 young, but fully expanded leaves from each plant. We scanned the leaves and quantified morphology using the morphometric software ‘*Lamina*’ (Bylesjo et al. 2008), from which we analysed four leaf traits: leaf area, leaf complexity 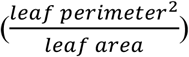, the number of leaf indents standardised by the perimeter and Specific Leaf Area 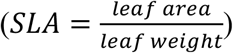. These leaf measurements represent leaf morphology and investment traits that show plastic responses to the abiotic environment in *S. chrysanthemifolius* (Walter et al. 2022; Walter et al. 2021), and in other plant systems, including sunflowers (Royer et al. 2009). We also used a Dualex instrument (Force-A, France) to measure the flavonol pigment content of the leaf. Flavonols are secondary metabolites that combat oxidative stress created by stressful abiotic (e.g. light and temperature) and biotic (e.g. herbivore) conditions (Mierziak et al. 2014).

### Quantifying genetic variance in fitness

To quantify genetic variance in fitness, we used *MCMCglmm* (Hadfield 2010) within R (v3.6.1; R Core Team 2019) to apply the generalised linear mixed effects model

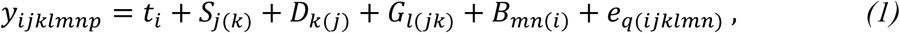

where the only fixed effect was transplant elevation (*t_i_*). The random effects *S*_*j*(*k*)_ represented the *j*th sire, *D*_*k*(*j*)_ the *k*th dam and *G*_*l*(*jk*)_ the *l*th individual of the breeding design nested within dam and sire. We included fitness as a univariate, poisson-distributed response variable (*y_ijklmnp_*), which quantifies genetic variance in relative fitness 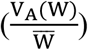 when estimates from a generalised linear model with a log-normal distribution are obtained on the latent scale (Bonnet et al. 2019; Morrissey and Bonnet 2019). To quantify genetic variance in relative fitness at each transplant elevation (and the genetic covariance among elevations), we estimated a 3×3 covariance matrix for the sire component by specifying random slopes and intercepts for transplant elevation.

To account for differences between transplant dates and among experimental blocks (within transplant date) at each elevation, we included the *n*th experimental block from the *p*th transplant date as a random effect (*b*_*mn*(*i*)_). Preliminary analyses showed that including separate random effects for transplant date and experimental block produced identical estimates of genetic variance. *e*_*q*(*ijklmn*)_ represents the error variance. We included a substantial burn-in and thinning interval to allow model convergence, which we confirmed by checking that effective sample sizes exceeded 85% of the number of saved samples. We used uninformative parameter-expanded priors for the random effects (Hadfield 2010), which we checked by changing the scale parameter and ensuring that there was no effect on the posterior distribution.

### Connecting genetic variance in plasticity with fitness

#### Identifying the ecological importance of leaf traits

For all traits in the subsequent analyses, we calculated the average across all leaves sampled from each plant. To test whether variation in leaf traits was associated with fitness at each elevation, we used multiple regression by applying generalised linear mixed effects models with ‘*lme4*’ (Bates et al. 2015) for each elevation. We included all five traits as continuous predictors (each standardised by their global mean) and the number of flowers as a poisson-distributed response variable. Experimental block and genotype were included as random effects.

We then used a multivariate analysis of variance to test whether plasticity significantly changed the multivariate phenotype across elevation by including all five traits as the multivariate response variable, transplant elevation as the categorical dependent variable, and the experimental blocks within transplant elevation as the error term.

#### Genetic correlations between leaf plasticity and fitness

We calculated plasticity across elevation by standardizing all clones of each genotype at 1,500m and 2,000 m by dividing by the mean value of that genotype at the home site (500 m). This standardization calculated the trait values for each of the 314 genotypes at the novel elevation, relative to their trait value at the home site (see **Fig. 4B-C**). We estimated plasticity separately for each transplant date. To quantify the genetic correlation between phenotypic plasticity and fitness, we used equation 1, but only for the data collected at 2,000 m and with all five leaf traits and fitness (number of flowers) as the multivariate response variable. This calculates the covariance between plasticity in each trait and fitness at 2,000 m. We extracted the sire component and calculated the genetic correlations between each trait and fitness.

**Fig. 3.**
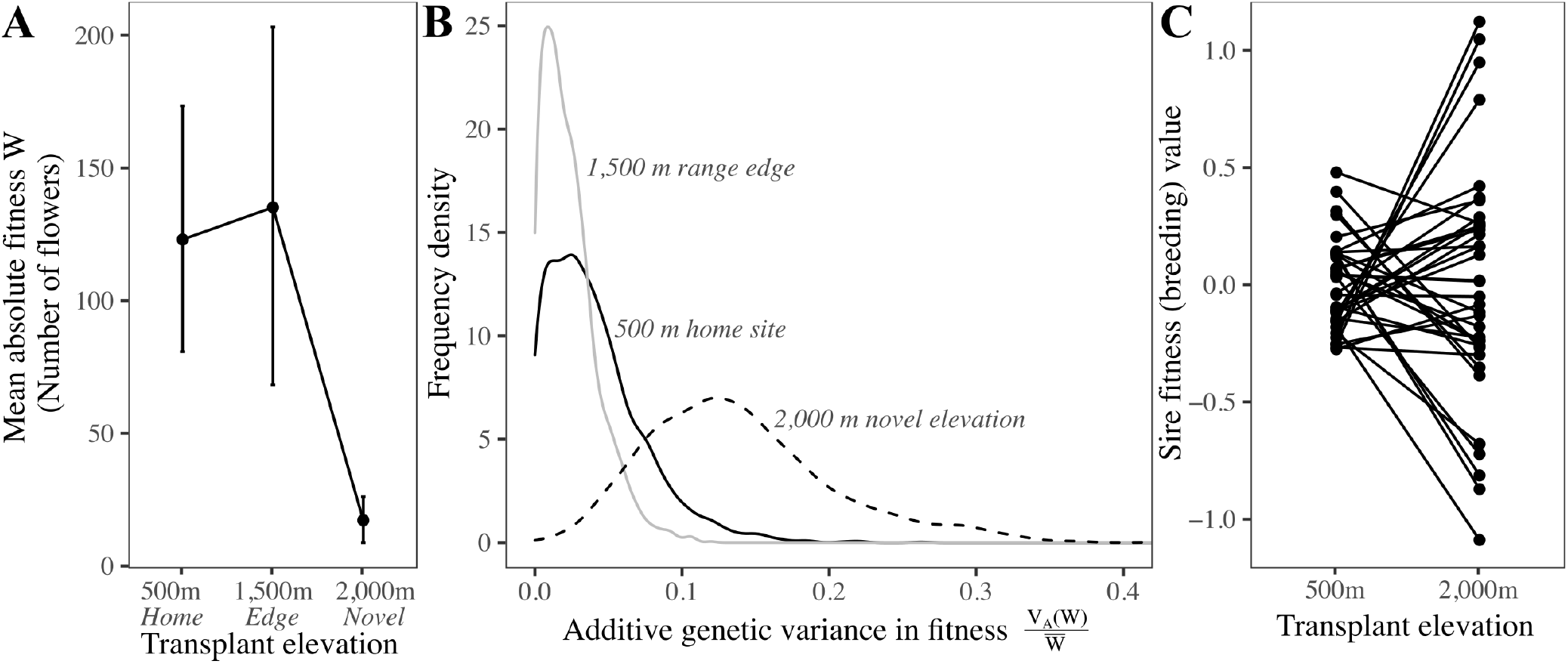
Absolute mean fitness 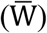 was 87% lower at the novel elevation, and this was associated with an increase in additive genetic variance in fitness 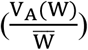. (**A**) Mean absolute fitness dropped outside the native range. Credible intervals represent 95% Highest Posterior Density (HPD) intervals of the mean. (**B**) Posterior distributions for the estimates of additive genetic variance in fitness. Genetic variance in fitness was significantly greater at the novel elevation (2,000 m) where distributions did not overlap at 90% HPD with the edge of the range, and 80% with the native site. (**C**) By visualising the fitness values for the sires (i.e. their breeding values), we can see how the genotypes respond differently to the native (500 m) and novel (2,000 m) elevations relative to the population mean (represented as zero). Reflecting the increase of genetic variance in fitness at the novel elevation (from **B**), differences among genotypes are greater at 2,000 m than at 500 m. The weak genetic correlation between the native and novel elevation (−0.1; −0.76, 0.49 HPD; **Table S2**) suggests that genotypes varied in the extent that they reduced fitness at the novel elevation. Genotypes that performed the best outside the range performed relatively poorly within the range, which is visualised by the high fitness genotypes at 2,000 m having low fitness at 500 m.

**Fig. 4.**
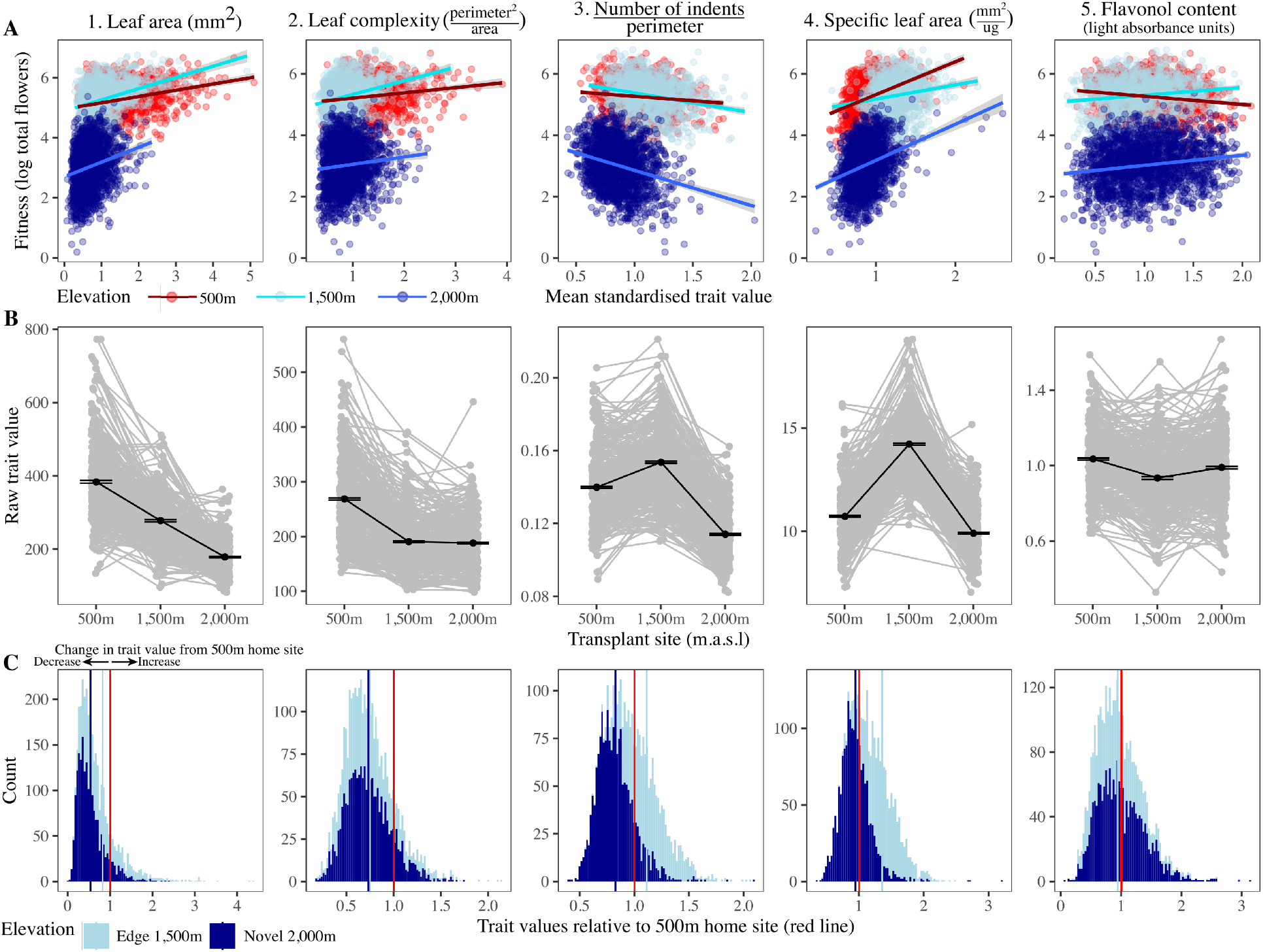
Changes in phenotype and selection across elevations. **(A)** Phenotype-fitness associations at each elevation. Trait values are standardised by the global mean for each trait. Each point represents an individual plant and shows that selection is in a similar direction for all elevations, except for flavonol content. (**B**) The change in raw trait values across elevation. Black lines and circles represent the overall mean (±1 SE), with all the genotypes from the breeding design in grey. Plasticity from the home site (500 m) to the 1,500m is in the same direction as the 2,000 m site for leaf area, complexity and flavonol content. For all traits, except specific leaf area, plasticity shows a stronger change in magnitude from 500 m to the novel 2,000 m elevation, when compared to the 1,500 m range edge. (**C**) Frequency distribution for plasticity in each trait, calculated as the degree to which each genotype changes trait values from the home site (vertical red line) to the edge of the range (light blue) and to the novel elevation (dark blue). Values of one represent no change from the home site, whereas values above and below one represent plasticity as an increase and decrease in the trait value, respectively.

### Gene expression analyses

To test whether gene expression variation across elevation was associated with fitness, logistical constraints made it necessary to conduct analyses on a subset of the 314 genotypes. We chose two sets of genotypes: The six genotypes that showed the greatest fitness at 2,000 m represent genotypes that increase the adaptive potential (‘*AP*’ genotypes) of the population at the novel elevation, and the six genotypes with the lowest fitness at 2,000 m that represent genotypes more specialised to conditions with their native range (‘*HR*’ genotypes for ‘*Home Range*’) (**Fig. S2A**). See **Methods S3** for details on how genotypes were chosen.

At each transplant elevation, we sampled leaves from three randomly selected clones of each of the 12 chosen genotypes. Plants were sampled when they were flowering but were still growing vegetatively. From each plant, we sampled 3-4 young leaves (c.15mm long), which we immediately submerged in RNAlater and stored at 4°C for 24 hours and then at −80°C before RNA extraction. To reduce environmental variation, we sampled on three consecutive days (19-21 November) between 9am and 11am under similar weather conditions.

We extracted RNA for the leaves of each plant and quantified gene expression at each elevation using 3’ RNAseq (QuantSeq). See **Methods S3** for RNA extraction protocols and transcriptome assembly. Once assembled, we annotated the transcriptome using *Trinotate* and identified orthologous genes from our transcriptome in *Arabidopsis thaliana* using *OrthoFinder* (Emms and Kelly 2019). 3’ reads were mapped to the reference transcriptome using Salmon v1.1.0 (Patro et al. 2017). 72.6-87.3% of reads were mapped. Transcript abundance estimates were imported into R using *txImport* (Soneson et al. 2015), with estimates normalised according to library size but not by transcript length, as the 3’ sequencing method removes this bias. We visualised the broad patterns of variation in gene expression by normalizing transcript abundance using the variance stabilizing transformation function with *DESeq2* (Love et al. 2014) followed by a principal components analysis **(Fig. S3**). We then calculated differential expression of transcripts using non-normalised transcript counts between transplant sites for each set of genotypes (*AP* and *HR*) separately using *DESeq2*. Differentially expressed genes were defined as those showing both significant (adjusted p-value <0.01) and strong (log2-fold change >2 or <-2) changes in expression. We then tested for enrichment of gene ontology (GO) terms for differentially expressed genes using a Kolmogorov-Smirnoff test and Fisher’s exact test with *topGO* (Alexa and Rahnenführer 2019).

## RESULTS

### (1) Reduced mean fitness but increased genetic variance in fitness in the novel environment

Consistent with our first prediction (**Fig. 2A**), absolute mean fitness (number of flowers) was high within the native range (500 m and 1,500 m) but decreased by 87% at 2,000 m (**Fig. 3A**). Also consistent with our first prediction, additive genetic variance in relative fitness 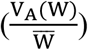 was near zero within the native range (500 m and 1,500 m), but more than three times greater at 2,000 m, which predicted a 14% increase in mean fitness in the subsequent generation (**Fig. 3B**; **Table S2**). Near zero genetic variance in fitness within the native range suggests that all genotypes maintained high fitness in familiar environments. By contrast, at the novel elevation genotypes failed (to differing extents) to maintain high fitness, increasing genetic variation in fitness.

We also found that genotypes with greater fitness at the novel elevation show lower fitness at the native site (**Fig. 3C**), as indicated by a weak and non-significant negative genetic correlation of −0.1 (−0.76, 0.49 90% Highest Posterior Density interval [HPD]) for fitness between the native site and the novel elevation. While a positive correlation would suggest that genotypes that perform well at one site also perform well at the other site, a [non-significant] correlation near zero suggests that genotypes respond differently by showing changes in relative fitness across elevations. The near-zero correlation in our results shows that while many genotypes do not show large changes in relative fitness across sites (**Fig. 3C**), those genotypes with the greatest fitness at 2,000 m have lower fitness relative to other genotypes in the native range. Genotypes that can increase the adaptive potential of the population in novel environments could therefore have a selective disadvantage within their native range.

The ten sires with the highest fitness in the novel environment were collected from three of the five sites (**Table S3**), suggesting that genetic variation from particular sites could be important for increasing the adaptive potential in novel environments. However, differences among sampling sites only accounted for a small proportion (0.5%) of the total variance in fitness compared to additive genetic variance (9.3%) (**Methods S4** and **Table S8**). These results, combined with no evidence of local adaptation among sites (**Methods S4** and **Fig. S7**), suggest that the five sampling sites are essentially all part of the same population.

### (2) Genetic variance in fitness in the novel environment correlates closely with plasticity

#### Identifying the ecological importance of leaf traits

We found that variance among genotypes was greater than among clones within genotype (**Methods S2**), suggesting that multiple clones provide a reliable representation of the response of each genotype to each elevation. We also observed a significant association between all leaf traits and fitness at each elevation (**Fig. 4A** and **Table S4**), suggesting that these leaf traits are important for maintaining fitness as the environment varies. At all transplant elevations, selection is in a similar direction, except for flavonol content, where greater flavonol content is associated with greater fitness at 1,500m and 2,000m, but with lower fitness at the 500 m home site (**Fig. 4A**).

In support of other transplant experiments that measured the same traits (Walter et al. 2022; Walter et al. 2021), we found evidence of plasticity as large changes in leaf traits across elevation (**Fig. 4B**) associated with a large and significant change in mean multivariate phenotype with elevation (MANOVA *F*_2,23_ = 44.521, *P*<0.0001). Plasticity reduced leaf area, leaf complexity and flavonol content similarly at the edge of the range (500-1,500 m) and the novel elevation (500-2,000 m). The number of indents and SLA increased from the home site to the edge of the range, but then decreased at the novel elevation. Only leaf area showed a greater magnitude of plastic change in phenotype at 2,000 m compared to 1,500 m (**Fig. 4B**).

While selection at 2,000 m favoured larger values of all traits, except the number of indents (**Fig. 4A**), plasticity created a reduction in trait values from the home site to the novel 2,000 m elevation (**Fig. 4B**), which suggests that fitness at 2,000 m is likely to be greater for genotypes that change their phenotype less across elevation.

#### Genetic correlations between leaf plasticity and fitness

To test whether the increased genetic variance in fitness at the novel elevation was associated with plasticity, we focussed on estimating genetic correlations between plasticity and fitness at 2,000 m. To quantify plasticity, we divided the trait values of each genotype at 2,000 m by the mean trait value of that genotype at the home site. This standardization calculates the trait values at 2,000 m relative to the home site for each genotype. Values of 1 reflect no change in phenotype between sites, while values above and below 1 respectively reflect increases and decreases in trait values from the home site (**Fig. 4C**). We predicted that if genetic differences in plasticity underlie the observed increased genetic variance in fitness at 2,000 m, we would observe significant genetic correlations between trait plasticity and fitness (**Fig. 2B**).

As predicted, we found strong and significant genetic correlations between plasticity and fitness at the novel elevation (>90% of the posterior distribution did not overlap with zero) for four traits: leaf area (0.47; 0.01, 0.88 90% HPD), the number of indents (−0.51; −0.95, −0.07 HPD), SLA (0.61; 0.27, 0.94 HPD) and flavonol content (0.55; 0.13, 0.95 HPD) (**Fig. 5A**). The increase in genetic variance in fitness at the novel elevation was therefore genetically correlated with plasticity for four of five leaf traits, suggesting that genetic variation in plasticity increases the adaptive potential of a population exposed to a novel environment. Only plasticity in leaf complexity showed no association with fitness (−0.03; −0.54, 0.53 HPD) (**Fig. 5A**).

**Fig. 5.**
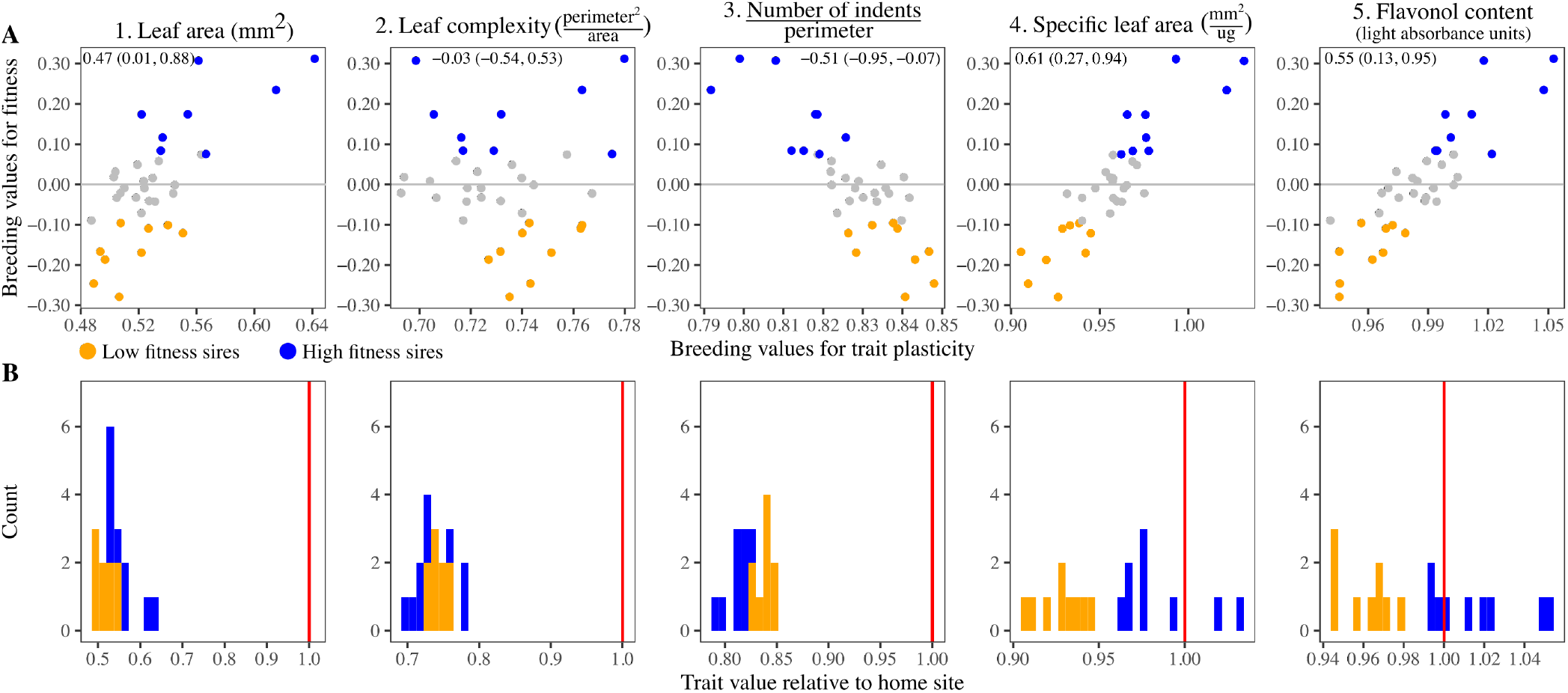
Genetic variance in plasticity was significantly correlated with fitness at the novel elevation for all traits, except leaf complexity. (**A**) Visualization of the genetic correlations using the 36 sire genetic (i.e. breeding) values for plasticity (x-axis) versus fitness (y-axis). Sire genotypes with the greatest fitness (blue circles) at 2,000 m show different levels of plasticity to genotypes with lowest fitness (orange circles). Inset text presents the posterior mean (and 90% HPD intervals) of the genetic correlations between plasticity and fitness at 2,000 m. (**B**) Histogram of the sire genetic values for trait plasticity that represent the magnitude of plastic change in trait means from the home site (value of 1, represented by the red vertical line) to 2,000 m. Compared to low fitness sires (orange), higher fitness at 2,000 m is associated with leaf plasticity as a smaller reduction in leaf area, SLA and flavonol content, but a larger reduction in the number of indents.

Genetic variation in fitness was associated with different patterns of plasticity that depended on the trait. For the number of indents, genotypes with higher fitness at 2,000 m show greater plasticity than low fitness genotypes, created by stronger reductions in trait values between 500 m and 2,000 m (**Fig. 5B)**. By contrast, for leaf area, SLA and flavonol content, genotypes with higher fitness at 2,000 m show lower plasticity across elevation than the genotypes with lower fitness (**Fig. 5B**). Reaction norms for the genotypes with the highest and lowest fitness at 2,000 m shows how plasticity across elevation is associated with fitness at the novel elevation. **Fig. 6A** shows that high fitness *AP* (*adaptive potential*) genotypes show different changes across elevation, and often arrive at different trait values at 2,000m when compared to the low fitness *HR* (*home range*) genotypes. Differences in how the leaves change across elevation for high (*AP*) and low (*HR*) fitness genotypes are visualised in **Fig. 6B**.

**Fig. 6.**
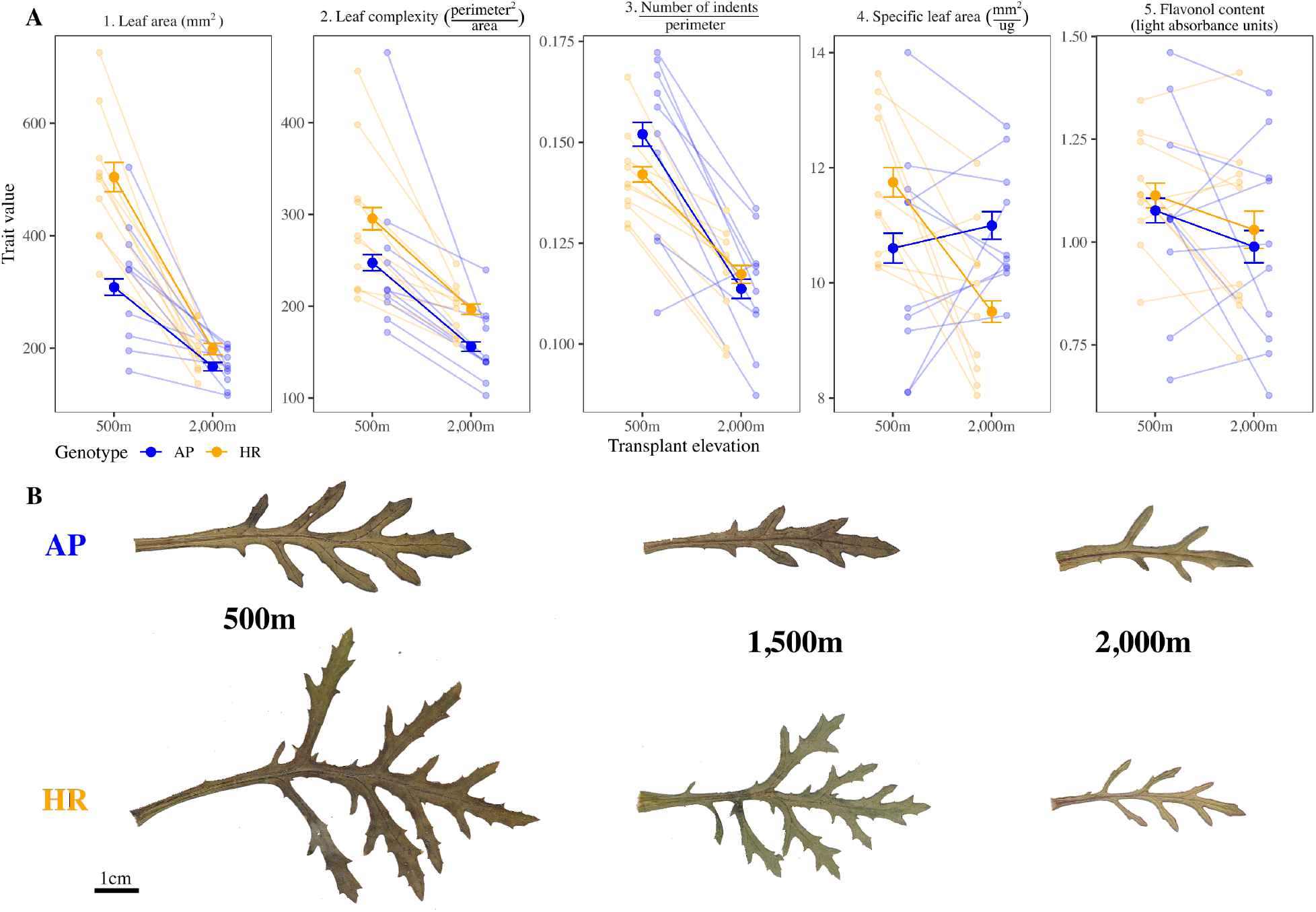
Changes in leaf traits across elevation for genotypes selected for the gene expression analysis. *AP* (*adaptive potential*) genotypes (blue) showed the higher fitness at 2,000 m, compared to the low fitness *HR* (*home range*) genotypes (orange). (**A**) Large circles with error bars (±1 SE) represent the average for the 10 *AP* and *HR* genotypes (represented by small circles). *AP* and *HR* genotypes show different patterns of plasticity for leaf area, number of indents and specific leaf area. Note that plasticity in flavonol content is not different for *HR* vs *AP* genotypes, which is because although fitness shows a strong genetic correlation with plasticity, the phenotypic correlation is weak. (**B**) Images of leaves for an *AP* (top row) and *HR* (bottom row) genotype across elevation. While both genotypes show reduced leaf area and leaf complexity at higher elevations, *AP* genotypes show less of a reduction in leaf area and more of a reduction in the number of leaf indents across elevations, when compared to *HR* genotypes. These changes are associated with differences between *AP* and *HR* in gene expression across elevation for genes relating to leaf development and morphogenesis (see gene expression results and **Fig. S5**).

### (3) Plasticity in gene expression is associated with fitness in the novel environment

To test whether differences in fitness at the novel elevation were associated with differences in gene expression, we sampled and analysed RNA from clones (at all three elevations) of genotypes that showed high and low fitness at 2,000 m. Overall, more genes were differentially expressed at 2,000 m (i.e., showed higher or lower expression levels relative to the native site) compared to the range edge, suggesting that plasticity under more novel conditions is created by broad transcriptional responses across the genome. *AP* (*adaptive potential*) genotypes with greater fitness at 2,000 m showed significant changes (adj. *P*<0.01) in more genes than the low fitness *HR* (*home range*) genotypes (1,376 in *AP* vs 514 genes in *HR*) within the range (500 m-1,500 m), suggesting that the *HR* genotypes adjusted fewer genes to maintain high fitness within their range (**Fig. 7A**). However, the number of genes that showed expression changes tripled outside the native range (500 m-2,000 m) for both classes of genotypes (*AP*=4,876 genes vs *HR*=4,720) **(Fig. 7B**; **Fig. S4**). The mean magnitude of expression change was also greater at the novel elevation (*AP*=1.41, *HR*=1.21) when compared to the edge of the range (*AP*=1.19, *HR*=1.04). Compared to the *HR* genotypes, *AP* genotypes showed strong overexpression in 10× more genes at the novel elevation, but underexpressed half as many genes (**Fig. 7C**). Therefore, as predicted (**Fig. 2C**), distinct patterns of gene expression between novel and native environments were associated with genotypic differences in fitness at the novel elevation.

**Fig. 7.**
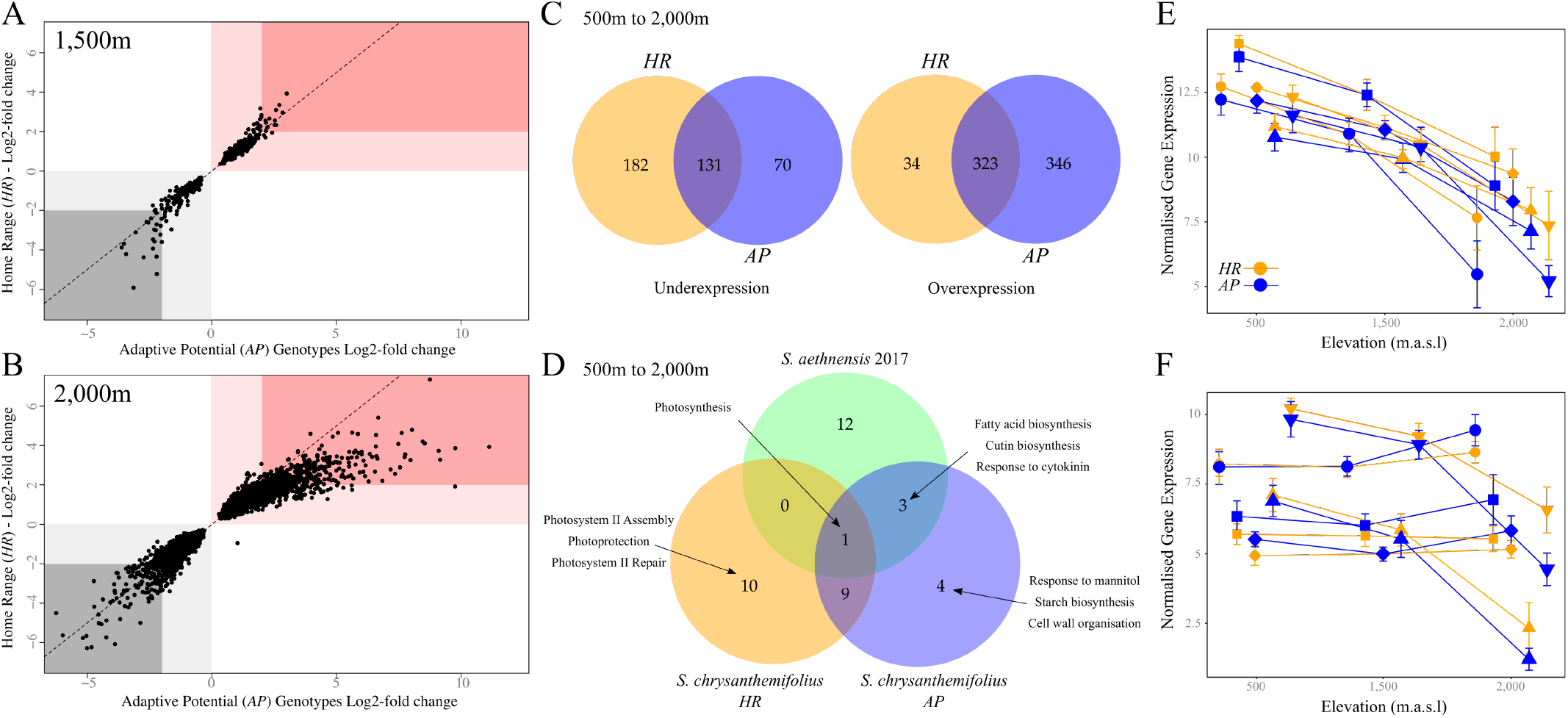
*AP* and *HR* genotypes differ in gene expression at the novel elevation. (**A-B**) Comparing expression of the same genes (black circles), with deviations from the dotted line representing differences in expression between *AP* (x-axis) versus *HR* (y-axis) genotypes. Red and grey shading represents overexpressed and underexpressed genes, respectively. Darker shading denotes strong expression changes. **(A)** Within the native range (1,500 m vs 500 m), more genes were differentially expressed in *AP* compared to *HR* genotypes. (**B**) Outside the native range (2,000 m vs 500 m), more genes were differentially expressed, and more genes in the *AP* genotypes showed a greater magnitude of differential gene expression. **(C)** Numbers of differentially expressed genes (adjusted *P*<0.01 and log-fold change <2 or >-2 for over- and underexpression) for the *AP* and *HR* genotypes between the home site (500 m) and outside the range (2,000 m). *AP* genotypes overexpressed more genes than *HR* genotypes. **(D)** Enriched Gene Ontology (GO) terms for differentially expressed genes for both genotypes and for a high elevation sister species (*S. aethnensis*) transplanted along the same elevation gradient in 2017 (Walter et al. 2022). Only the three most significant terms are shown. *AP* genotypes shared more GO terms with *S. aethnensis*, suggesting that high fitness at the high elevation is associated with similar genetic pathways to the native species. **(E)** Comparing elevational changes in mean expression for five genes (associated with light harvesting) for the *AP* (blue) and *HR* (orange) genotypes. Each gene is represented by a different shape and credible intervals represent 95% confidence intervals. *AP* genotypes showed stronger underexpression at 2,000 m. **(F)** Elevational changes in mean expression for five genes associated with responses to cold. *AP* genotypes show stronger under- and overexpression, but the reaction was gene-dependent.

#### Gene ontology

We also predicted that greater fitness at 2,000 m would be associated with changes in gene expression for ecologically-important genes. *AP* and *HR* genotypes differed in the functional categories of genes that varied in expression at the novel elevation (**Fig. 7D**). *AP* genotypes showed differential expression of genes relating to biosynthesis, whereas *HR* genotypes differentially expressed genes related to photosystems. Comparing the enriched GO terms to those of a 2017 transplant experiment (Walter et al. 2022) that included the high elevation sister species (*S. aethnensis*), we found that four (of 17) GO terms were enriched in *S. aethnensis* as well as in the *AP* genotypes, compared to only one (of 20) GO term shared by *S. aethnensis* and the *HR* genotypes (**Fig. 7D**). Compared to the *HR* genotypes, *AP* genotypes of *S. chrysanthemifolius* therefore showed a more similar gene expression response to the closely-related *Senecio* species native to the 2,000 m habitat.

GO terms that were significantly enriched at 2,000 m included light harvesting in photosystem I, cell wall organization, and responses to cold (**Table S5**). To illustrate the contrasting patterns of expression between *AP* and *HR* genotypes, we selected the five most differentially expressed genes from each GO term and compared the reaction of these genotypes across elevation. For genes involved in photosystem I (**Fig. 7E**), *AP* genotypes showed stronger underexpression than *HR* genotypes at the novel elevation. For genes involved in cold responses **(Fig. 7F)**, *AP* genotypes showed a stronger response than *HR* genotypes, which included greater overexpression or underexpression depending on the gene.

We then tested whether differences in gene expression across elevations for *AP* and *HR* genotypes were associated with genes known to be associated with leaf development and morphogenesis in *Arabidopsis* (Huala et al. 2001). We found that 86 *Arabidopsis* orthologs were differentially expressed in either *AP* or *HR* genotypes between 500 and 2,000 m (**Table S6**). These included two genes that are important for determining leaf shape and dissection in *Arabidopsis*: PINFORMED1 (PIN1) and ASYMMETRIC LEAVES1 (AS1) (Barkoulas et al. 2008). In both cases, the orthologous genes in *S. chrysanthemifolius* showed significantly lower expression at 2,000 m, with a greater decrease in expression shown by *AP* genotypes **(Fig. S5)**, which supports the finding that greater reductions in leaf indentation is associated with higher fitness at 2,000 m (**Fig. 6**).

## DISCUSSION

We provide empirical support for two fundamental hypotheses that are crucial for understanding how populations respond to novel environments: First, we provide strong evidence that although adaptive plasticity fails in novel environments and causes declines in absolute mean fitness (**Fig. 3A**), as predicted by theory, genetic variance in fitness increases and improves the adaptive potential of the population, which could allow rapid adaptation to the novel environment (**Fig. 3B**). Second, increased genetic variance in fitness in the novel environment was associated with plasticity as elevational changes in leaf traits (**Fig. 6**), and as changes in gene expression in genes important for responding to the novel high-elevation habitat (**Fig. 7**), which suggests that genetic variation in plasticity increases the adaptive potential of populations exposed to novel environments.

Previous studies have shown high levels of genetic variance for fitness in natural populations (Hendry et al. 2018; Kulbaba et al. 2019; Sheth et al. 2018), heritable variation in plasticity (Nussey et al. 2005), adaptive plasticity in phenology (Charmantier et al. 2008) and rapid evolutionary responses to range shifts (Buckley and Bridle 2014). However, to our knowledge, this study provides the first experimental evidence that genetic variation in plasticity is associated with increased genetic variance in [a trait closely-associated with] fitness in a novel environment, which increases the adaptive potential of the population in a novel environment. Although these results suggest that rapid adaptation to the novel environment could occur when beneficial alleles rapidly increase in frequency, population persistence via evolutionary rescue will only be likely if the population size in the novel environment remains large enough so as to avoid extinction (Bridle et al. 2019; Chevin et al. 2013; Gonzalez et al. 2013; Polechová and Barton 2015), as discussed in detail below. However, the increased genetic variance in fitness in the novel environment that we observed suggests that genetic variation important for rapid adaptation is already segregating in the population, which means that evolutionary rescue will be more likely than if adaptation were to rely on new mutation (Orr and Unckless 2014).

Our results also support evidence that genetic variance in fitness depends on the environment in which it is quantified (Kulbaba et al. 2019; Sheth et al. 2018), and show for the first time that genotypes suitable for adaptation to novel environments may be under negative selection in native environments. Genotypes with higher fitness at the novel environment tended to show lower relative fitness in the native environment (**Fig. 3C**). Genotypes important for evolutionary rescue may therefore be selected against in native environments (perhaps at early life history stages). If so, then identifying genetic variation important for evolutionary rescue will be difficult to identify within the native range because these genotypes are maintained at low frequency throughout much of a species’ range (Brennan et al. 2019). As shown in our results, adaptive plasticity is also likely to hide genetic variation important for evolutionary rescue because differences in plasticity among genotypes are much smaller and difficult to detect within the native range. Our results therefore suggest that studies estimating genetic variation in fitness *in situ* are likely to underestimate the potential for evolutionary rescue because alleles that increase adaptive potential in novel environments are only seen in those environments, and are hidden by adaptive plasticity and selection within the native range.

### The role of genetic variance in plasticity for increasing adaptive potential in novel environments

Genetic variance in fitness at the novel environment was associated with distinct patterns of plasticity in leaf traits and gene expression. Compared to low fitness genotypes at the novel elevation, high fitness genotypes were associated with greater plasticity as larger reductions in leaf indentation, but lower plasticity that caused smaller reductions in leaf area, and the maintenance of trait values across elevations for specific leaf area and flavonol content (**Fig. 5–6**). Similarly, our gene expression data suggest that the potential to adapt to novel environments is driven by genotypes with the most beneficial gene expression profiles (Josephs 2021; Wang and Althoff 2019). Compared to the low fitness *HR* genotypes, high fitness *AP* genotypes differentially expressed more genes at 2,000 m, which was associated with many genes exhibiting stronger overexpression (**Fig. 7C**). The changes observed in leaf morphology across elevations were supported by changes in expression for genes that regulate leaf development and morphogenesis (AS1 and PN1). Differential expression in the high fitness genotypes occurred in functional categories associated with plasticity in the closely related high elevation sister species. Together, these results suggest that population persistence in novel environments will require genotypes that possess adaptive plasticity of a particular magnitude and direction (Chevin and Hoffmann 2017; Lande 2009).

Given that genetic differences in fitness were associated with distinct patterns of plasticity, our findings contrast with evidence that adaptation to novel environments involves non-adaptive plasticity in gene expression for Trinidadian guppies responding to predators (Ghalambor et al. 2015), or that a lack of genetic variance in gene expression will prevent beneficial responses to novel environments in butterflies exposed to seasonal fluctuations (Oostra et al. 2018). In our results, genotypes with high fitness at the novel environment showed greater changes in gene expression but smaller changes in three of five leaf traits, which suggests that greater gene expression plasticity could reduce the amount of plasticity in leaf traits across elevations, and that this ‘phenotypic homeostasis’ may help to prevent larger reductions in fitness (Velotta and Cheviron 2018). Such phenotypic homeostasis could therefore be critical for adaptation to novel environments given that the direction of plasticity is unlikely to be adaptive in novel environments, and that greater plasticity will be more likely to reduce than increase fitness (Hoffmann and Bridle 2021).

We previously found that at the novel 2,000 m elevation, plasticity moved the phenotype of *S. chrysanthemifolius* towards the native phenotype of its close relative, *S. aethnensis* (Walter et al. 2022), suggesting that plasticity in *S. chrysanthemifolius* is, to some extent, adaptive at 2,000 m. In the current study, high fitness was maintained at the edge of the range, which was associated with plasticity that involved large changes in phenotype and gene expression. At 2,000m, genotypes of *S. chrysanthemifolius* with greater fitness displayed similar patterns of gene expression to *S. aethnensis* with responses occurring in genes important for responding to elevation. High fitness genotypes also showed smaller reductions in specific leaf area and flavonol than low fitness genotypes, which is in the direction of the native phenotype of *S. aethnensis*. Furthermore, leaf complexity was lower at higher elevations, which is also in the direction of the native phenotype. Together, these results provide strong evidence that plasticity in *S. chrysanthemifolius* is adaptive to the edge of the range and, to some extent, in the novel 2,000 m habitat. It is therefore likely that for plasticity to help the population persist in a novel environment while adaptation occurs, plasticity will need to be at least partly adaptive (Ghalambor et al. 2007).

### Greater adaptive potential does not necessarily lead to evolutionary rescue

The observed increase in genetic variance in fitness at the 2,000 m site suggests that genetic variation segregating in natural populations can promote adaptation to the novel elevation. However, the likelihood of persistence in novel environments will depend on whether the population size is maintained at level that allows adaptation to remain viable (Bridle et al. 2019; Chevin et al. 2013; Gonzalez et al. 2013; Polechová and Barton 2015). In particular, where a population experiences a large decline in mean fitness, evolutionary rescue may be unlikely even if suitable genetic variation is available to selection. This is because the effective size of the population is likely to be too small to allow selection to overcome drift, making extinction more likely than adaptation (Bridle and Vines 2007; Carlson et al. 2014; Chevin et al. 2013). It is possible that adaptive plasticity can help the population persist while adaptation occurs by allowing a greater number of individuals to survive in the novel environment (Ghalambor et al. 2007), but we require further data to understand when and where adaptive plasticity fails to maintain viable population sizes. Empirical estimates of ecological and life history parameters are therefore needed to identify when and where population declines are not so strong as to prevent evolutionary rescue (Connallon and Sgrò 2018; Hoffmann and Bridle 2021).

Our results suggest that evolutionary rescue in response to ongoing environmental change will depend on selection on genetic variation in plastic responses (Chevin and Hoffmann 2017; Chevin and Lande 2011; Lande 2009), which would fine-tune gene-products and trait values in the novel environments. Genetic variation for sensitivity to the environment is likely to be unevenly distributed across a species’ range (Colautti and Barrett 2013; Sheth and Angert 2016; Stratton 1994; Walter et al. 2020), with local adaptation, mutation and drift likely to determine which populations contain the genetic variation that can increase the potential for evolutionary rescue (Hargreaves and Eckert 2019; Hoffmann and Bridle 2021). To understand when and where rapid adaptation will allow the persistence of populations exposed to novel environmental regimes, genetic variation in environmental sensitivity needs to be estimated for multiple populations exposed to a range of novel environments.

## Acknowledgements

We thank David Aguirre, Spencer Barrett, Roger Butlin, Robert Dugand, Ary Hoffmann and Carla Sgrò for advice and comments on previous versions. We are very grateful to those who helped with fieldwork: Alessandro Barbato, Octavia Brayley, Guy Burstein, Maria Castrogiovanni, Stefania Catara, Sarah du Plessis, Carmen Impelluso, Enrico la Spina, Mari Majorana, Jessica Menzies, Morgan Millen, Giuseppe Pepe and Daniel Ward. We thank Piante Faro for providing glasshouse resources.

## Funding

This work was supported by NERC grants NE/P001793/1 and NE/P002145/1 awarded to JB and SH.

## Author contributions

GW, SH and JB designed the experiment with input from SC, JC and AC. GW, AC and DT conducted the experiments. JC collected, extracted and processed RNA samples, and analyzed the transcriptome data. GW analyzed the fitness and phenotype data. GW wrote the manuscript with JC, JB and SH, and input from all other authors.

## Declaration of interests

The authors declare no competing interests.

## Notes

### Competing Interest Statement

The authors have declared no competing interest.

### Summary of Updates

Revised text and updated figures

