## Supplementary material for "Hidden genetic variation in plasticity provides the potential for rapid adaptation to novel environments"

**This PDF file includes:**

**Table S1** Sampling locations

**Fig. S1** Map of experiment and sampling locations

**Fig. S2** Genotype choice for gene expression analysis

**Fig. S3** Visualizing variation in gene expression across genotypes, clones and elevation

**Table S2** Additive genetic variance matrix for fitness across elevation

**Table S3** Sire breeding values for fitness at 2,000 m

**Table S4** Multiple regression to quantify phenotype-fitness associations

**Fig. S4** Histogram of differentially expressed genes

**Table S5** Significant GO terms for differentially expressed genes

**Table S6** Differentially expressed genes for leaf development and shape in *Arabidopsis*

**Fig. S5** Expression changes in the genes that function in *Arabidopsis* leaf morphology

**Methods S1** Comparing fitness as the number of flowers versus seed production

**Methods S2** Relating phenotypic traits to the elevational gradient

**Methods S3** RNA extraction, sequencing and transcriptome assembly

**Methods S4** Contribution of site variance to estimates of genetic variance

**Table S1:** Location of sampled individuals used as the parental generation for the crossing design used in the glasshouse. See **Fig. S1** for a map of locations.

| Site | Elevation | Latitude | Longitude | # Sires | # Dams |
|---|---|---|---|---|---|
| Bonnano | 790 m | 37°38'24.92"N | 15° 2'50.80"E | 9 | 10 |
| Cacciola | 680 m | 37°37'31.32"N | 15° 3'26.71"E | 6 | 6 |
| Poggofelice | 590 m | 37°39'44.31"N | 15° 5'48.55"E | 6 | 5 |
| Spina | 730 m | 37°39'19.27"N | 15° 4'30.92"E | 9 | 10 |
| Trecastagni | 570 m | 37°36'46.67"N | 15° 4'29.64"E | 6 | 5 |
| **Totals** | | | | **36** | **36** |

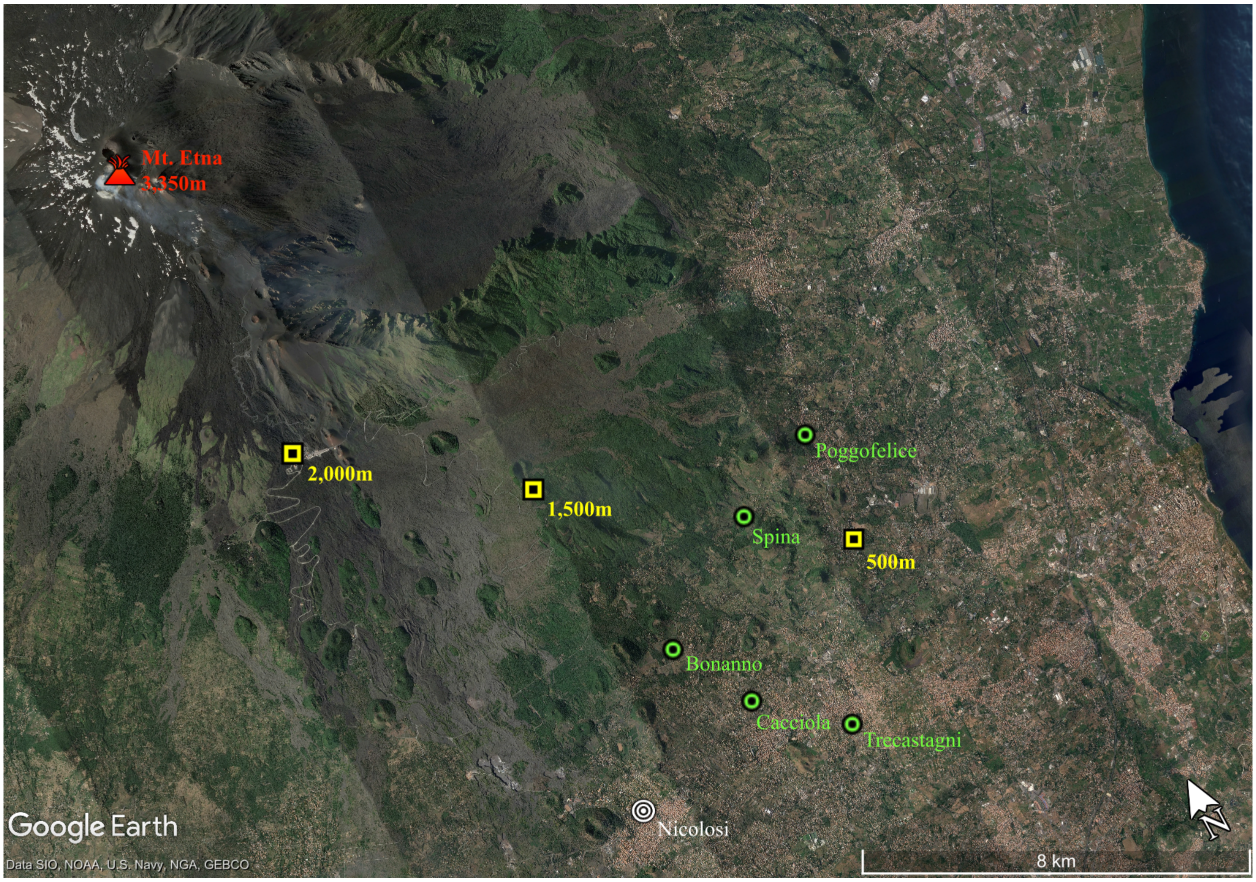


**Fig. S1** Map of the experiment. Transplant sites (yellow squares) lie along a south-eastern transect. Sites where genotypes for the parental generation were sampled are represented by green circles. See **Table S1** for coordinates.

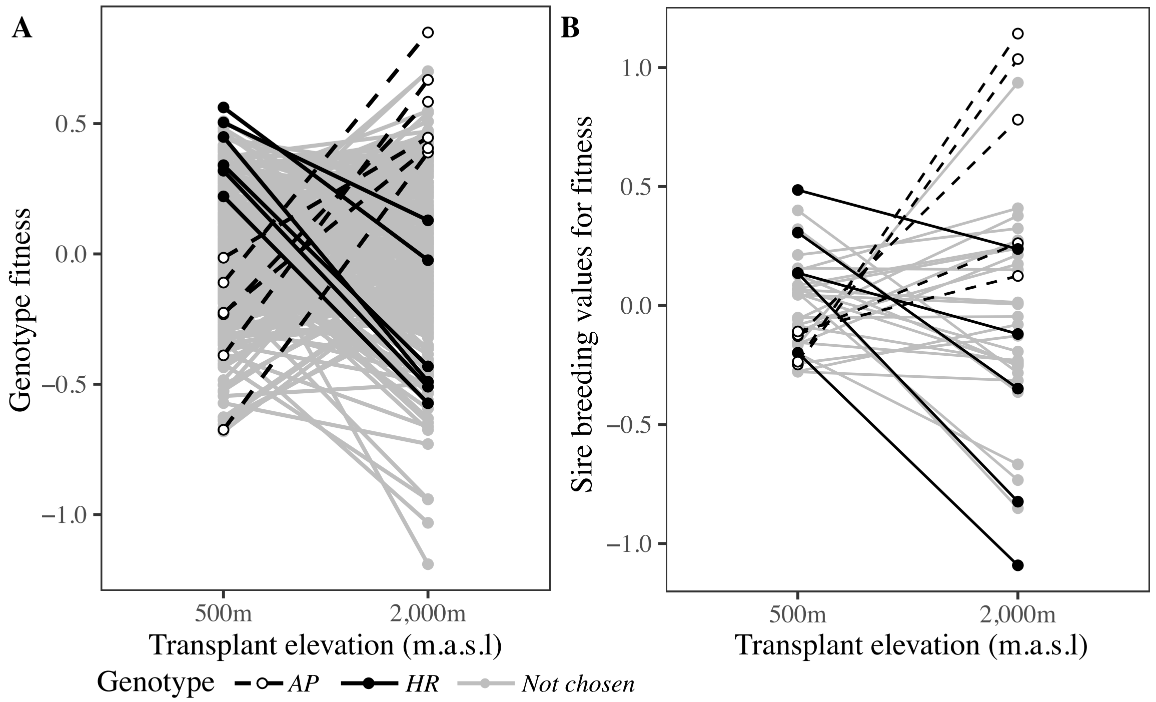


**Fig. S2** Selection of genotypes based on their fitness response between elevational extremes, calculated using equation 1. (**A)** Chosen genotypes were based on changes in relative fitness from the home site (500 m) to outside the range (2,000 m) for all offspring of the crossing design. Unfilled circles and broken lines represent the *AP* genotypes, and filled circles and solid lines represent the *HR* genotypes, that were chosen for the gene expression analysis. Gray lines and circles represent the remaining genotypes from the crossing design that were not chosen. (**B)** The 12 genotypes chosen for the gene expression analysis were from 10 sires that also showed large changes in relative fitness. Therefore, genotypes chosen for the study of differential expression also represented additive genetic variance in fitness.

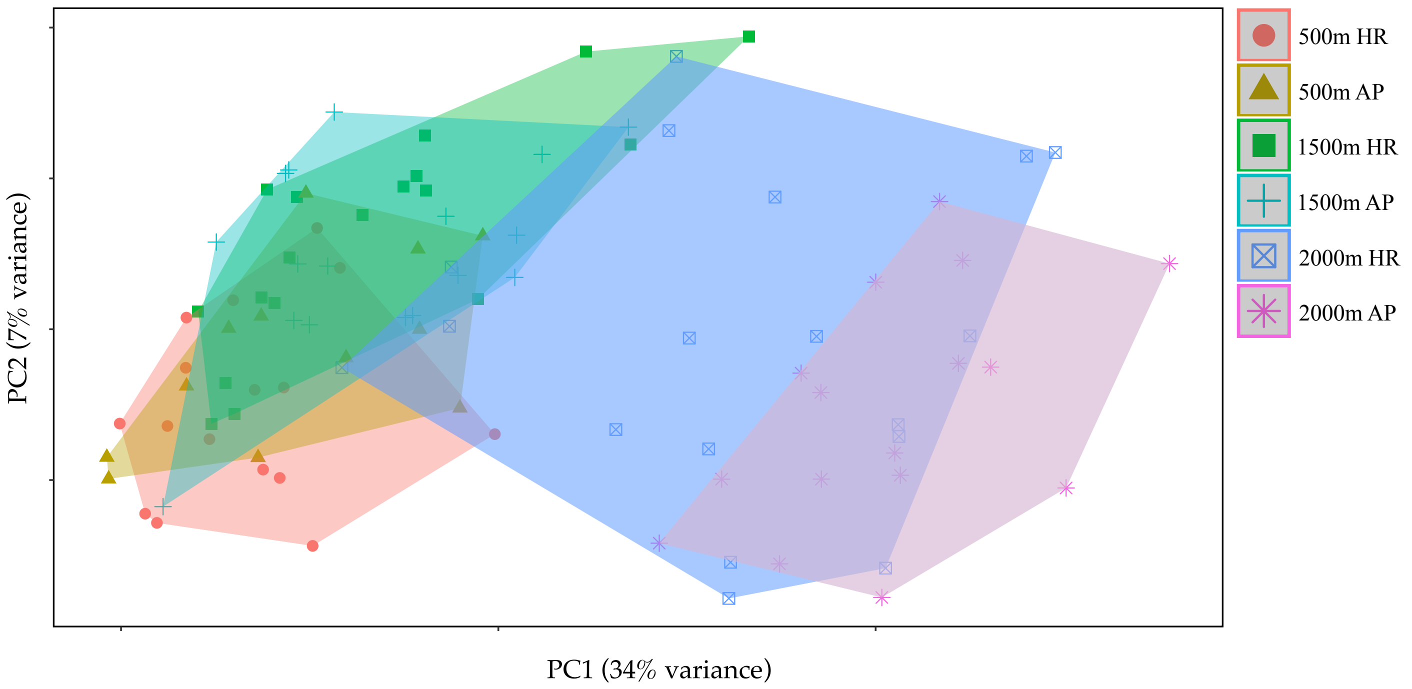


**Fig. S3** Variation in gene expression among the 98 samples. Individuals represent clones belonging to one of 12 genotypes (6 *AP* and 6 *HR*) and are colored according to transplant site (meters above sea level) and genotype (*AP* and *HR*).

**Table S2**: The additive genetic covariance matrix for fitness. The diagonal contains the additive genetic variance in fitness at each elevation. Genetic covariances between elevations are presented above the diagonal, and the genetic correlations between elevations are presented below the diagonal. Numbers in parentheses denote 90% HPD intervals.

| | 500 m | 1,500 m | 2,000 m |
|---|---|---|---|
| 500 m | 0.041 (0,0.084) | 0.007 (-0.012,0.027) | -0.006 (-0.047,0.036) |
| 1,500 m | 0.18 (-0.42,0.88) | 0.024 (0,0.05) | 0.019 (-0.01,0.055) |
| 2,000 m | -0.1 (-0.76,0.49) | 0.35 (-0.18,0.86) | 0.14 (0.041,0.24) |

**Table S3** Breeding values for each sire arranged in order of fitness at 2,000 m. Colors represent the different source sites from where the parental sires were sampled.

| Site | Sire | BLUP for fitness at 2,000 m |
|---|---|---|
| Trecastagni | 43 | -1.0888447 |
| Poggofelice | 38 | -0.8715609 |
| Trecastagni | 7 | -0.813119 |
| Cacciola | 67 | -0.7222338 |
| Poggofelice | 19 | -0.6791872 |
| Trecastagni | 62 | -0.3871309 |
| Poggofelice | 26 | -0.3526613 |
| Trecastagni | 49 | -0.2990977 |
| Cacciola | 51 | -0.2676762 |
| Poggofelice | 32 | -0.2585369 |
| Bonnano | 44 | -0.2393247 |
| Spina | 15 | -0.2377734 |
| Spina | 56 | -0.222774 |
| Cacciola | 31 | -0.1794271 |
| Spina | 45 | -0.1316659 |
| Cacciola | 21 | -0.1168628 |
| Spina | 50 | -0.0840763 |
| Trecastagni | 69 | -0.0503502 |
| Cacciola | 14 | 0.01473255 |
| Spina | 8 | 0.01862107 |
| Spina | 68 | 0.1276029 |
| Bonnano | 33 | 0.16324835 |
| Spina | 2 | 0.16472873 |
| Poggofelice | 61 | 0.21531416 |
| Bonnano | 20 | 0.23538244 |
| Bonnano | 37 | 0.24987902 |
| Bonnano | 9 | 0.2545542 |
| Poggofelice | 57 | 0.26270505 |
| Bonnano | 39 | 0.28823247 |
| Bonnano | 13 | 0.3598893 |
| Cacciola | 1 | 0.37181873 |
| Bonnano | 25 | 0.42108935 |
| Trecastagni | 3 | 0.78915086 |
| Bonnano | 55 | 0.94840234 |
| Spina | 27 | 1.04734513 |
| Spina | 63 | 1.12295376 |

**Table S4** Summary statistics for the multiple regressions applied to each transplant site. $R^2$ values are provided for the fixed effects alone (marginal) and when taking into account both fixed and random effects that include environmental block and genotype within the crossing design (conditional). Statistical tests for significant association between traits and fitness were conducted using log-likelihood ratio tests ($\chi^2$ = Chi-square statistic for the likelihood ratio test), which show that leaf traits were significantly associated with fitness at all transplant elevations.

| **Elevation** | **Trait** | **Esti-mate** | **95% CI** | $\chi^2$ | **P-value** |
|---|---|---|---|---|---|
| 500 m | Intercept | 5.245 | 5.027, 5.463 | | |
| Marginal $R^2$=10.08% | 1. Leaf area ($mm^2$) | 0.014 | 0.006, 0.022 | 11.58 | 0.00067 |
| | 2. Leaf complexity ($perimeter^2$/area) | 0.137 | 0.121, 0.153 | 280.41 | <0.0001 |
| Conditional $R^2$=98.33% | 3. Number of indents (# / perimeter) | -0.609 | -0.644, -0.574 | 1166.61 | <0.0001 |
| | 4. SLA ($mm^2$ / µg) | 0.545 | 0.515, 0.574 | 1327.72 | <0.0001 |
| | 5. Flavonol content (light absorbance) | -0.152 | -0.168, -0.136 | 360 | <0.0001 |
| 1,500 m | Intercept | 4.927 | 4.479, 5.376 | | |
| Marginal $R^2$=6.86% | 1. Leaf area | 0.119 | 0.108, 0.131 | 402.62 | <0.0001 |
| | 2. Leaf complexity | 0.255 | 0.235, 0.276 | 597.24 | <0.0001 |
| Conditional $R^2$=98.95% | 3. Number of indents | -0.312 | -0.341, -0.284 | 458.1 | <0.0001 |
| | 4. SLA | 0.143 | 0.118, 0.168 | 123.28 | <0.0001 |
| | 5. Flavonol content | 0.115 | 0.1, 0.13 | 221.45 | <0.0001 |
| 2,000 m | Intercept | 2.388 | 1.987, 2.789 | | |
| Marginal $R^2$=15.37% | 1. Leaf area | 0.286 | 0.238, 0.334 | 134.33 | <0.0001 |
| | 2. Leaf complexity | 0.23 | 0.18, 0.279 | 81.39 | <0.0001 |
| Conditional $R^2$=94.67% | 3. Number of indents | -0.745 | -0.825, -0.666 | 345.94 | <0.0001 |
| | 4. SLA | 0.939 | 0.88, 0.999 | 943.24 | <0.0001 |
| | 5. Flavonol content | 0.159 | 0.125, 0.193 | 83.44 | <0.0001 |

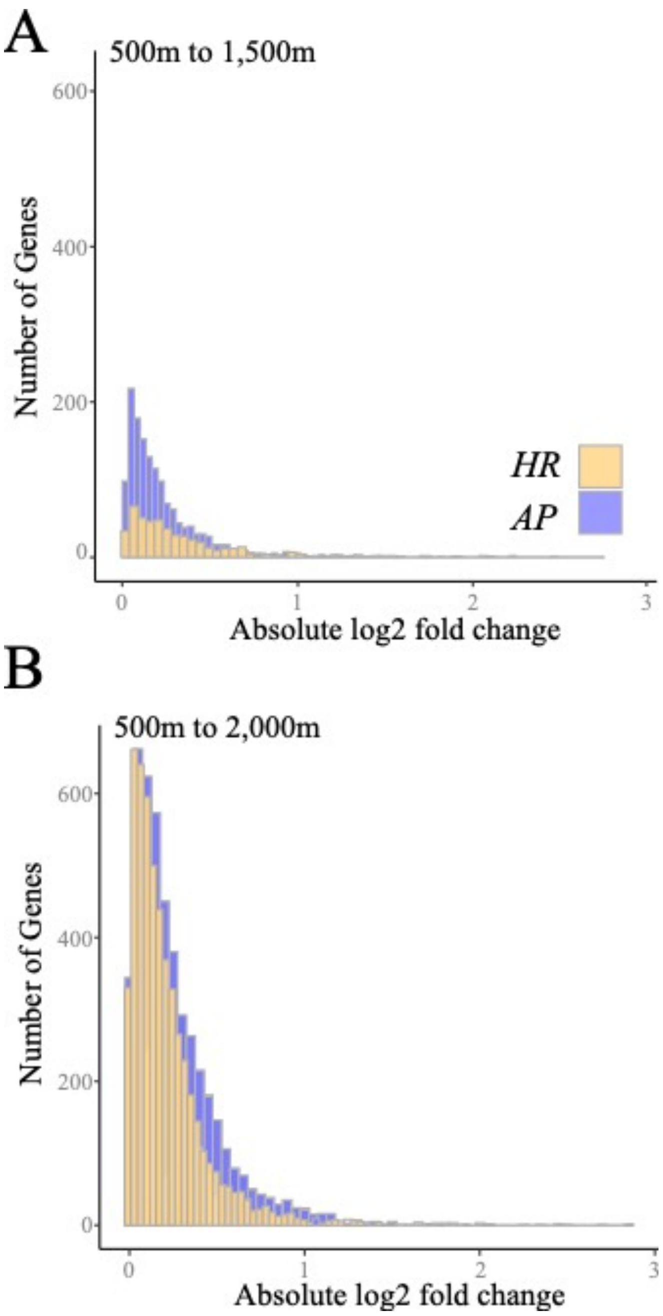


**Fig. S4** The strength of differential expression for *HR* (orange) vs *AP* (blue) genes. (**A**) Within the native range (500-1,500 m), *AP* genotypes differentially express more genes compared to *HR* genotypes. (**B**) Outside the native range (500-2,000 m), both genotypes differentially express more genes, with *AP* genotypes showing greater magnitude of expression changes.

**Table S5** Significantly enriched GO terms between the two sets of genotypes (*AP* and *HR*) at 2,000 m. Significance defined as p-value < 0.05 following both a Kolmogorov Smirnoff (KS) test and Fisher's exact test (Fisher's).

| *Gene Ontology Term* | *Gene Ontology Description* | p-value (KS) | p-value (Fisher's) |
|---|---|---|---|
| GO:0009765 | Photosynthesis, light harvesting | 7.6e-7 | 1.3e-8 |
| GO:0018298 | Protein-chromophore linkage | 1.6e-6 | 2.2e-6 |
| GO:0009768 | Photosynthesis, light harvesting in PSI | 3.9e-6 | 0.0006 |
| GO:0071555 | Cell wall organization | 2.5e-6 | 0.0018 |
| GO:0019684 | Photosynthesis, light reaction | 0.0005 | 1.2e-5 |
| GO:0010143 | Cutin biosynthetic process | 0.001 | 0.0008 |
| GO:0009809 | Lignin biosynthetic process | 0.003 | 0.03 |
| GO:0009409 | Response to cold | 0.004 | 0.0008 |
| GO:0010166 | Wax metabolic process | 0.005 | 0.02 |
| GO:0009831 | Plant-type cell wall modification | 0.007 | 0.01 |
| GO:0042335 | Cuticle development | 0.008 | 1.2e-5 |
| GO:0006949 | Syncitium formation | 0.009 | 0.02 |
| GO:0009807 | Lignan biosynthetic process | 0.014 | 0.006 |
| GO:0045493 | Xylan catabolism | 0.024 | 0.003 |
| GO:0009644 | Response to high light intensity | 0.024 | 0.007 |
| GO:0015976 | Carbon utilisation | 0.025 | 0.004 |

**Table S6.** Genes identified from the *Arabidopsis* TAIR database tagged with the keywords "Leaf development", "Leaf morphogenesis", "Leaf shape", "Leaf margin" and "Leaf lamina" that were differentially expressed in *AP* and *HR* genotypes between 500 and 2,000m.

| | | | | Adj. p-value (500 vs 2000m) | |
|---|---|---|---|---|---|
| TAIR locus | Name | Keyword | *S. chrys. locus* | AP | HR |
| AT3G22200 | POP2 | Leaf development | DN13080_c2_g1_i11 | 1.5E-06 | 9.8E-12 |
| | | | DN13080_c2_g2_i5 | 4.0E-02 | - |
| AT4G02570 | AXR6 | Leaf development | DN1738_c0_g1_i1 | 1.0E-02 | 9.4E-03 |
| | | | DN3264_c0_g1_i1 | 2.0E-02 | 3.6E-04 |
| AT1G53310 | PEPC1 | Leaf development | DN1761_c0_g1_i3 | 5.0E-10 | 1.6E-10 |
| | | | DN1761_c0_g1_i7 | 2.7E-10 | 7.4E-09 |
| AT2G37860 | RE | Leaf development | DN5928_c0_g1_i3 | 6.4E-03 | 2.0E-04 |
| AT1G14280 | PKS2 | Leaf development | DN3650_c0_g1_i2 | 6.9E-08 | - |
| AT2G42200 | SPL9 | Leaf development | DN12076_c0_g1_i2 | 2.0E-02 | 2.0E-03 |
| AT3G15030 | TCP4 | Leaf development | DN1499_c0_g1_i1 | 1.8E-02 | - |
| AT2G32280 | VCC | Leaf development | DN10816_c0_g1_i1 | 2.8E-03 | 4.8E-02 |
| AT1G53230 | TCP3 | Leaf development | DN1499_c0_g1_i1 | 1.8E-02 | - |
| AT1G10670 | ACLA-1 | Leaf development | DN14043_c0_g1_i6 | 5.6E-08 | 3.7E-12 |
| | | | DN3501_c0_g1_i1 | 9.1E-07 | 1.0E-03 |
| AT2G31070 | TCP10 | Leaf development | DN1499_c0_g1_i1 | 1.8E-02 | - |
| AT4G39400 | BIN1 | Leaf development | DN18034_c3_g1_i3 | 3.0E-02 | 3.1E-02 |
| AT2G42600 | PPC2 | Leaf development | DN1761_c0_g1_i3 | 5.4E-05 | 1.6E-10 |
| | | | DN1761_c0_g1_i7 | 2.7E-10 | 7.4E-09 |
| AT3G11450 | ZRF1A | Leaf development | DN2759_c0_g1_i1 | 1.5E-03 | 2.5E-02 |
| AT5G28640 | AN3 | Leaf development | DN4110_c0_g3_i1 | 2.9E-02 | - |
| AT5G16780 | DOT2 | Leaf development | DN17075_c0_g1_i4 | 3.3E-02 | - |
| AT3G08640 | RER3 | Leaf development | DN7745_c0_g1_i1 | - | 3.3E-02 |
| | | | DN3791_c2_g1_i2 | 4.7E-02 | 8.7E-04 |
| AT5G05620 | TUBG2 | Leaf development | DN18798_c0_g4_i1 | 2.7E-05 | 6.3E-05 |
| AT5G56030 | HSP81.2 | Leaf development | DN686_c0_g2_i2 | 2.7E-12 | 2.2E-02 |
| | | | DN686_c0_g3_i1 | 4.3E-05 | 6.0E-04 |
| AT5G53660 | GRF7 | Leaf development | DN13936_c2_g1_i1 | 5.2E-06 | 2.5E-07 |
| AT2G28350 | ARF10 | Leaf development | DN17022_c2_g1_i1 | 4.1E-05 | 8.3E-06 |
| | | | DN17200_c0_g1_i1 | 3.4E-02 | 3.7E-02 |
| | | | DN17353_c0_g1_i1 | 6.1E-05 | 4.2E-07 |
| AT5G04810 | PPR4 | Leaf development | DN8240_c0_g1_i1 | - | 2.2E-05 |
| AT1G08410 | DIG6 | Leaf development | DN2443_c0_g1_i1 | 4.2E-05 | 2.6E-02 |

| | | | | | |
|---|---|---|---|---|---|
| AT1G17980 | PAPS1 | Leaf development | DN863_c1_g1_i4 | 5.6E-06 | 6.1E-08 |
| AT5G10270 | CDKC1 | Leaf development | DN14557_c1_g2_i1 | 2.8E-03 | 4.4E-02 |
| AT5G64960 | CDKC2 | Leaf development | DN14557_c1_g2_i1 | 2.8E-03 | 4.4E-02 |
| AT3G14940 | PPC3 | Leaf development | DN1761_c0_g1_i3 | 5.4E-05 | 1.6E-10 |
| | | | DN1761_c0_g1_i7 | 2.7E-10 | 7.4E-09 |
| AT1G48920 | NUC-L1 | Leaf development | DN20546_c0_g1_i33 | 4.5E-07 | 1.4E-02 |
| | | | DN20776_c1_g1_i2 | 2.1E-04 | - |
| | | | DN670_c0_g1_i10 | 8.7E-03 | - |
| AT1G13260 | EDF4 | Leaf development | DN13187_c0_g2_i1 | 3.0E-02 | 2.4E-04 |
| AT4G33950 | OST1 | Leaf development | DN16117_c0_g3_i8 | 2.2E-02 | 2.6E-04 |
| AT4G31160 | DCAF1 | Leaf development | DN3273_c0_g2_i2 | 5.9E-04 | 3.5E-02 |
| AT4G00850 | GIF3 | Leaf development | DN5057_c0_g1_i1 | 1.1E-03 | 2.0E-06 |
| AT3G61650 | TUBG1 | Leaf development | DN18798_c0_g4_i1 | 2.7E-05 | 6.3E-05 |
| AT1G01160 | GIF2 | Leaf development | DN5057_c0_g1_i1 | 1.0E-03 | 2.0E-06 |
| AT2G16800 | CGF2 | Leaf development | DN14886_c4_g2_i1 | 3.0E-02 | - |
| AT2G28890 | PLL4 | Leaf development | DN72_c1_g1_i3 | 3.3E-05 | 5.9E-03 |
| AT1G70560 | CKRC1 | Leaf development | DN17357_c0_g1_i9 | 5.0E-05 | 1.1E-05 |
| AT4G20360 | SVR11 | Leaf development | DN18489_c1_g2_i1 | 1.7E-03 | 1.4E-11 |
| AT1G15690 | AVP1 | Leaf development | DN853_c1_g1_i3 | 1.6E-03 | 2.3E-07 |
| AT5G58230 | MSI1 | Leaf development | DN18640_c0_g2_i1 | 5.0E-02 | - |
| | | | DN8951_c0_g1_i1 | - | 1.7E-02 |
| AT4G02440 | EID1 | Leaf development | DN7960_c0_g1_i1 | 1.3E-03 | 8.2E-06 |
| AT1G07630 | PLL5 | Leaf development | DN72_c1_g1_i3 | 3.3E-05 | 5.9E-03 |
| AT4G30340 | DGK7 | Leaf development | DN3511_c0_g1_i4 | 2.1E-04 | - |
| AT4G15900 | PRL1 | Leaf development | DN15335_c1_g3_i2 | 1.0E-03 | 1.2E-02 |
| AT4G37650 | EAL1 | Leaf development | DN15500_c1_g2_i1 | 3.0E-03 | 4.0E-03 |
| AT1G56180 | VIR3 | Leaf development | DN17010_c0_g1_i3 | 8.9E-03 | - |
| AT3G15380 | CTL1 | Leaf development | DN405_c1_g1_i1 | 6.4E-04 | 1.6E-05 |
| AT2G40300 | FER4 | Leaf development | DN1831_c0_g1_i1 | 7.5E-08 | 4.3E-08 |
| AT1G79440 | ENF1 | Leaf development | DN3220_c0_g2_i1 | 2.0E-02 | - |
| AT4G24560 | UBP16 | Leaf development | DN1421_c0_g2_i3 | 2.2E-07 | 1.7E-07 |
| AT1G73590 | PIN1 | Leaf shape | DN17601_c0_g2_i3 | 9.3E-05 | 7.7E-04 |
| AT3G15730 | PLD | Leaf shape | DN6193_c0_g1_i6 | 1.0E-02 | - |
| AT2G34960 | CAT5 | Leaf margin | DN17698_c1_g1_i1 | 2.8E-02 | 3.2E-06 |
| | | | DN3644_c0_g1_i1 | 1.2E-02 | - |
| AT2G28680 | RmlC-like | Leaf margin | DN19627_c0_g1_i7 | 1.5E-07 | 2.3E-13 |
| AT2G39450 | MTP11 | Leaf margin | DN17749_c0_g1_i10 | 2.2E-02 | - |
| AT1G70560 | CKRC1 | Leaf margin | DN17357_c0_g1_i9 | 5.0E-05 | 1.1E-05 |
| AT1G52150 | CAN | Leaf morphogenesis | DN229_c0_g2_i1 | 1.9E-03 | 6.2E-05 |
| | | | DN229_c0_g3_i1 | - | 3.0E-02 |

| | | | | | |
|---|---|---|---|---|---|
| | | | DN3600_c0_g1_i3 | - | 3.0E-02 |
| AT5G39740 | RPL5B | Leaf morphogenesis | DN17570_c0_g1_i2 | 1.0E-02 | 5.5E-04 |
| AT3G15030 | TCP4 | Leaf morphogenesis | DN1499_c0_g1_i1 | 1.8E-02 | - |
| AT1G55250 | HUB2 | Leaf morphogenesis | DN1008_c0_g1_i1 | 6.6E-03 | 4.0E-04 |
| AT2G37630 | AS1 | Leaf morphogenesis | DN9323_c0_g1_i1 | 2.0E-07 | 2.0E-03 |
| AT3G05040 | HST1 | Leaf morphogenesis | DN3921_c0_g1_i2 | 3.0E-02 | - |
| AT1G48410 | AGO1 | Leaf morphogenesis | DN1568_c2_g3_i1 | - | 2.0E-02 |
| AT1G53230 | TCP3 | Leaf morphogenesis | DN1499_c0_g1_i1 | 1.8E-02 | - |
| AT2G23760 | SAW2 | Leaf morphogenesis | DN15861_c0_g2_i2 | 6.4E-06 | 2.2E-09 |
| | | | DN6003_c0_g1_i1 | 8.4E-09 | 2.0E-03 |
| AT3G25520 | RPL5A | Leaf morphogenesis | DN17570_c0_g1_i2 | 1.1E-02 | 5.0E-04 |
| | | | DN17570_c0_g1_i5 | | 1.4E-02 |
| AT2G31070 | TCP10 | Leaf morphogenesis | DN1499_c0_g1_i1 | 1.0E-02 | - |
| AT4G36870 | SAW1 | Leaf morphogenesis | DN15861_c0_g2_i2 | 6.4E-06 | 2.2E-09 |
| | | | DN6003_c0_g1_i1 | 8.4E-09 | 1.9E-03 |
| AT4G34740 | ASE2 | Leaf morphogenesis | DN6101_c0_g1_i1 | - | 2.0E-03 |
| AT2G17040 | NAC36 | Leaf morphogenesis | DN8530_c0_g1_i1 | - | 4.5E-03 |
| AT3G20630 | UBP14 | Leaf morphogenesis | DN1956_c0_g1_i2 | 3.0E-04 | 2.0E-03 |
| AT1G14400 | UBC1 | Leaf morphogenesis | DN15508_c1_g1_i2 | 2.0E-02 | - |
| AT1G01510 | AN | Leaf morphogenesis | DN2861_c0_g1_i1 | 2.0E-02 | - |
| AT4G00100 | PFL2 | Leaf morphogenesis | DN1650_c0_g1_i1 | 3.0E-03 | 9.0E-03 |
| AT5G08370 | AGAL2 | Leaf morphogenesis | DN2943_c0_g1_i2 | 5.0E-03 | 6.0E-03 |
| AT4G03550 | EED3 | Leaf morphogenesis | DN0_c0_g1_i2 | 8.0E-03 | 1.2E-05 |
| AT3G53020 | RPL24 | Leaf morphogenesis | DN17975_c0_g1_i1 | - | 4.0E-02 |
| AT4G29040 | RPT2A | Leaf morphogenesis | DN13563_c1_g1_i3 | 1.0E-02 | 1.0E-02 |
| AT1G26440 | UPS5 | Leaf lamina | DN490_c0_g1_i12 | 8.5E-07 | 9.1E-06 |
| AT2G03530 | UPS2 | Leaf lamina | DN490_c0_g1_i12 | 8.5E-07 | 9.1E-06 |
| AT2G47220 | DUF5 | Leaf lamina | DN19824_c2_g1_i4 | 4.0E-03 | - |
| AT2G26540 | DUF3 | Leaf lamina | DN13523_c0_g1_i1 | 5.0E-03 | 6.6E-04 |
| AT5G08000 | PDCB2 | Leaf lamina | DN15838_c0_g4_i1 | 8.5E-25 | 2.9E-16 |
| AT5G58787 | IRP4 | Leaf lamina | DN14981_c4_g1_i1 | 1.1E-02 | 6.9E-07 |
| | | | DN17802_c2_g1_i11 | 8.0E-03 | 1.8E-02 |
| AT5G61130 | PDCB1 | Leaf lamina | DN15838_c0_g4_i1 | 8.5E-25 | 2.9E-16 |
| AT5G38030 | DTX30 | Leaf lamina | DN6970_c0_g2_i1 | 2.0E-02 | - |

81

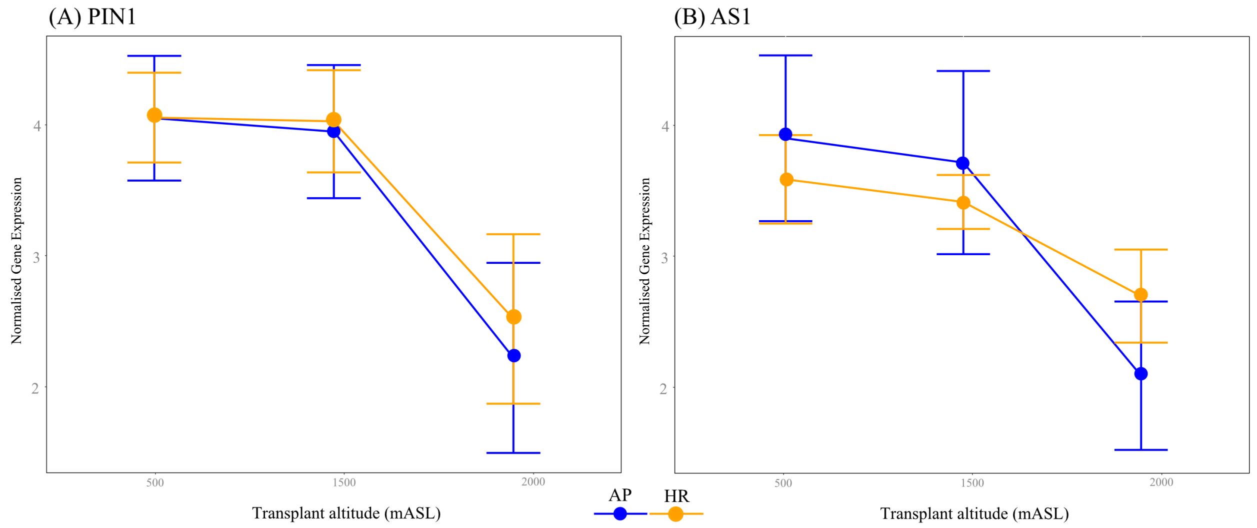


**Fig. S5** Comparing elevational changes in mean expression for the *AP* (blue) and *HR* (orange) genotypes orthologs for two genes of known function in *Arabidopsis:* **(A)** PIN1 gene that functions in leaf development; and **(B)** AS1 gene that functions in leaf shape. The mean expression for each gene is represented by a circle and credible intervals represent 95% confidence intervals.

***Methods S1. Comparing fitness as the number of flowers versus seed production***

In a previous transplant (Walter *et al.*, 2022), we compared the number of flowers to the proportion of seeds produced. For randomly selected individuals transplanted at the elevational extremes (500 m $n$ = 28; 2,000 m $n$ = 40), we counted the number of flowers and then collected mature seed heads on two sampling dates. Viable seeds were considered those that were large, brown and round; unviable seeds were those that were thin, empty and generally white. We used a linear mixed effects model implemented with '*lme4*' (Bates *et al.*, 2015) to test whether the number of flowers was associated with seed set. We included seed set as the response variable, and transplant site and number of flowers (and their interaction) as the fixed effects. Block within transplant site was the only random effect. We predicted that if plants that produced a large number of flowers also produced a large number of seed, then we would observe no association between the number of flowers produced and the average seed set per flower. This would suggest that the number of flowers provides a good estimate of total fitness because plants that produce more flowers would also produce more seeds. We found that the interaction between the number of flowers and transplant site was not significant ($\chi^2(1) = 0.7856$, $P$ = 0.3754), suggesting that the association between flower number and seed production was high and consistent across elevation (**Fig. S6**).

We then used the Kenwood-Roger approach, which is used to test the significance of fixed effects, to test whether the number of flowers was associated with seed set. As predicted, we found no significant effect of the number of flowers on average seeds produced per flower ($F_{1,27.1}$ = 0.2089, P = 0.6513), indicating that the number of seeds produced per inflorescence was similar for all plants. The number of flowers therefore provides a good representation for the total number of seeds produced for each plant, and provides a robust estimate of total fitness.

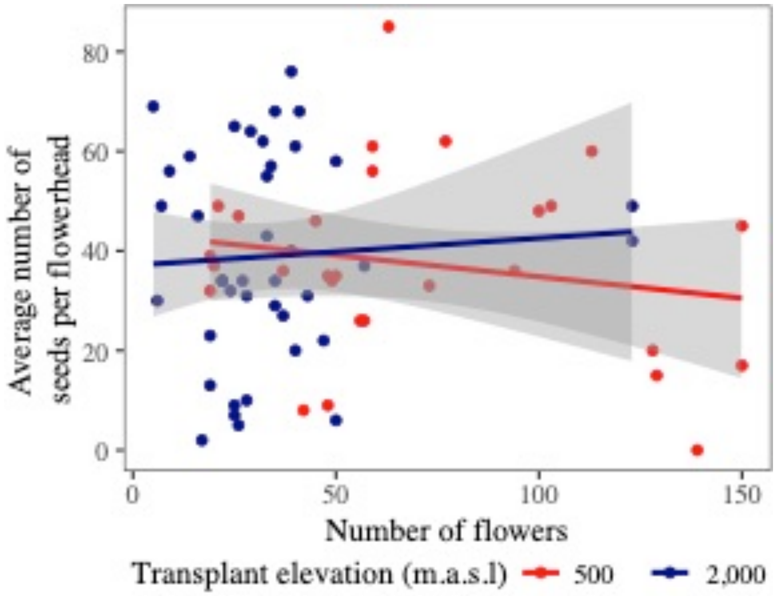


**Fig. S6** The number of flowers produced by a plant was not correlated with the number of seeds produced, and this was consistent for both transplant sites. This suggests that the number of flowers correlates with the number of seeds.

***Methods S2. Relating phenotypic traits to the elevational gradient***

Phenotypic variances (variance among all plants) tended to decrease with altitude, except for flavonol content, which increased in variance (**Table S7**). Using analysis of variance on each trait independently, we tested whether differences among the 314 genotypes in the breeding design described more variance than among clones within genotypes. We also included transplant site and experimental block nested within transplant site. All traits showed greater variance among genotypes than among clones within genotypes (leaf area: $F_{313,4592} = 2.963$, P<0.0001; leaf complexity: $F_{313,4592} = 11.761$, P<0.0001; number of indents: $F_{313,4592} = 8.962$, P<0.0001; SLA: $F_{313,4592} = 4.731$, P<0.0001; Flavonol content: $F_{313,4745} = 3.851$, P<0.0001. This meant that differences among the 314 genotypes accounted for >20% of the total variance in each trait (leaf area = 35.8%, leaf complexity = 51.3%, number of indents = 54.7%, SLA = 51.1% and flavonol content = 20.1%). Therefore, differences among genotypes were significant and multiple clones provided a reliable representation of the response of each genotype to environmental variation across elevation.

**Table S7** Total phenotypic variances (variances among all clones) for each trait at each elevation.

| | Area | Leaf complexity | Number of indents | SLA | Flavonol |
|---|---|---|---|---|---|
| 500 m | 0.255 | 0.125 | 0.032 | 0.045 | 0.082 |
| 1,500 m | 0.145 | 0.080 | 0.041 | 0.048 | 0.102 |
| 2,000 m | 0.050 | 0.063 | 0.027 | 0.040 | 0.124 |

***Methods S3. RNA extraction, sequencing and transcriptome assembly***

*Choosing genotypes*

Because our collection of young leaves for RNA samples had to be completed before the winter snow arrived, we were restricted to choosing genotypes before the flowers of all cuttings were counted. After flowers were counted for two clones of each genotype at each transplant site, we estimated genetic variation for fitness as the among-genotype variance (i.e. among the individuals in the crossing design) at each transplant site, and the covariance between transplants sites (see equation 1). We then chose the 15 genotypes with the highest (*AP* genotypes) and the 15 genotypes with the lowest (*HR* genotypes) relative fitness at 2,000 m. After fitness was quantified for all cuttings, we repeated the analysis for choosing the genotypes (this time with all the fitness data included), and from the sampled genotypes we chose the six *AP* genotypes and six *HR* genotypes for gene expression analysis that maintained the strongest fitness differences at 2,000 m (**Fig. S2A**). Importantly, chosen genotypes also represented changes in genetic variance in fitness, whereby sires of the chosen genotypes exhibited similar fitness responses to elevation (**Fig. S2B**).

*RNA extraction and sequencing*

For each clone, we homogenized all collected tissue and performed RNA extraction using a Qiagen RNeasy kit with β-mercaptoethanol added to the lysis buffer and included a DNase digestion step. We measured RNA purity and concentration using a Nanodrop ND1000 spectrophotometer and Qubit fluorometer. All sequencing and library preparation was performed at the Oxford Genomics Centre, The Wellcome Centre for Human Genetics, Oxford (UK). For each selected genotype, total RNA from a single individual was sequenced to produce 150bp reads using an Illumina NovaSeq6000 platform. For each individual, a small region close to the

3’ end of each transcript was sequenced to produce 75bp reads using a Lexogen QuantSeq library preparation and Illumina NextSeq500 platform (Moll *et al.*, 2014).

*Reference Transcriptome Assembly*

RNAseq of total RNA produced on average 28.75 million reads per sample. The 3’ sequencing produced on average 5.46 million reads per sample. Only samples that produced more than 1 million reads (n = 98) were included in downstream analyses. Quality assessment and trimming of all reads was performed using TrimGalore v0.6 (Phred quality cut-off = 20). Trimmed reads from the total RNAseq for each genotype were combined and a single reference transcriptome was *de novo* assembled in Trinity v2.8.4 (Haas *et al.*, 2013). To reduce transcript and isoform redundancy, we filtered the transcriptome using the EvidentialGene pipeline (minimum sequence length = 400 nucleotides) (Gilbert, 2019) and contaminating sequences were removed using the MCSC Decontamination pipeline (filter = Viridiplantae) (Lafond-Lapalme *et al.*, 2017). The reference transcriptome contained 29,224 transcripts with an n50 of 2,030 nucleotides. The reference transcriptome was annotated using the  pipeline (Bryant *et al.*, 2017). Nucleotide sequences were used to perform a *Diamond blastx* search and translated amino acid sequences were used to perform a *Diamond blastp* search, each of the *UniProt* database with a $1\times10^{-20}$ cut-off. This resulted in 22,335 unique annotations, and on average 1.06 annotations per transcript.

***Methods S4. Contribution of site variance to estimates of genetic variance***

Our estimates of genetic variance are taken from a population randomly sampled across five sites. If sites were to be very different, either due to local adaptation or genetic drift, then our estimates of genetic variance could be conflated by differences among sites. Although the interpretation of genetic variance with respect to the species remains the same because our estimates of genetic variance represent the adaptive potential for the randomly sampled parental generation from the broader population.

To ensure that site differences did not conflate our estimates of genetic variance we took two approaches: First, we tested for local adaptation using fitness data (collected using the same methods as the current study) from a previous experiment that transplanted individuals from all five sites at the same elevations (Walter *et al.*, 2022). We found no evidence of local adaptation (**Fig. S7**), suggesting that the sites have not undergone adaptive divergence within their home range.

Second, we re-analysed the fitness data from equation 1 in the main text, but included an additional random effects term ($c_p$) that estimates the variance among crosses and includes crosses conducted within sites (e.g. Spina×Spina) and among sites (e.g. Spina×Bonnano). We found that differences among sampling sites only accounted for a small proportion (0.5%) of the total variance in fitness compared to additive genetic variance estimated from the sire component (9.3%) (**Table S8**). These results, combined with no evidence of local adaptation among sites (**Fig. S7**), suggest that the five sampling sites are part of the same inter-connected population.

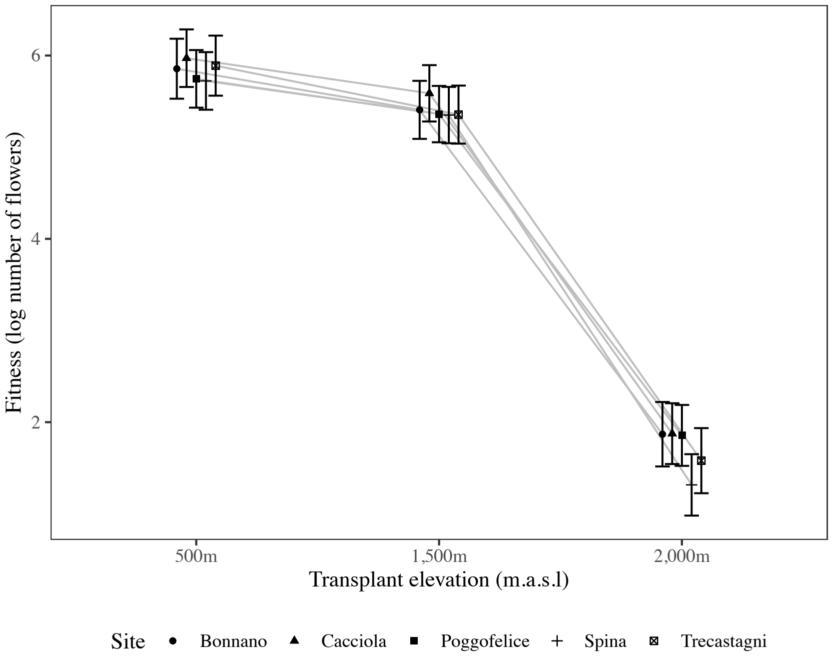


**Fig. S7** Fitness was similar for the genotypes of all five sites when transplanted as cuttings in 2019. This suggests that the five sites are not locally adapted and that they represent part of the same population. The experiment is described in detail in Walter *et al.* (2022).

**Table S8** Estimates of variance for each random component in equation 1, but also including the extra 'Site' component. Additive genetic variance is estimated as four times the sire variance.

| **Component** | **500 m** | **1,500 m** | **2,000 m** |
|---|---|---|---|
| Sire | 0.008 | 0.005 | 0.031 |
| Dam | 0.015 | 0.003 | 0.012 |
| Genotype | 0.078 | 0.051 | 0.111 |
| Site | 0.007 | 0.002 | 0.007 |
| Block | 0.371 | 0.467 | 0.554 |
| Residual | 0.545 | 0.379 | 0.614 |

***Supplementary references***


**Bates D, Machler M, Bolker BM, Walker SC. 2015.** Fitting linear mixed-effects models using lme4. *Journal of Statistical Software* **67**(1): 1-48.

**Bryant DM, Johnson K, DiTommaso T, Tickle T, Cougar MB, Payzin-Dogru D, Lee TJ, Leigh ND, Kuo TH, Davis FG, et al. 2017.** A tissue-mapped Axolotl de novo transcriptome enables identification of limb regeneration factors. *Cell Reports* **18**(3): 762-776.

**Gilbert DG. 2019.** Longest protein, longest transcript or most expression, for accurate gene reconstruction of transcriptomes? *bioRxiv*: 829184.

**Haas BJ, Papanicolaou A, Yassour M, Grabherr M, Blood PD, Bowden J, Couger MB, Eccles D, Li B, Lieber M. 2013.** De novo transcript sequence reconstruction from RNA-seq using the Trinity platform for reference generation and analysis. *Nature protocols* **8**(8): 1494.

**Lafond-Lapalme J, Duceppe M-O, Wang S, Moffett P, Mimee B. 2017.** A new method for decontamination of de novo transcriptomes using a hierarchical clustering algorithm. *Bioinformatics* **33**(9): 1293-1300.

**Moll P, Ante M, Seitz A, Reda T. 2014.** QuantSeq 3′ mRNA sequencing for RNA quantification. *Nature Methods* **11**(12): 972.

**Walter GM, Clark J, Cristaudo A, Nevado B, Catara S, Paunov M, Velikova V, Filatov D, Cozzolino S, Hiscock SJ, et al. 2022.** Adaptive divergence generates distinct plastic responses in two closely related *Senecio* species. *Evolution* **76**(6): 1229-1245.
